# Synthesis and degradation of FtsZ determines the first cell division in starved bacteria

**DOI:** 10.1101/314922

**Authors:** Karthik Sekar, Roberto Rusconi, Tobias Fuhrer, Elad Noor, Jen Nguyen, Vicente I. Fernandez, Marieke F. Buffing, Michael Berney, Roman Stocker, Uwe Sauer

## Abstract

In natural environments, microbes are typically non-dividing. Such quiescent cells manage fleeting nutrients and gauge when intra- and extracellular resources permit division. Quantitative prediction of the division event as a function of nutritional status is currently achieved through phenomenological models for nutrient-rich, exponentially growing cultures. Such models, however, cannot predict the first division of cells under limiting nutrient availability. To address this, we analyzed the metabolic capability of starved *Escherichia coli* that were fed pulsed glucose at defined frequencies. Real-time metabolomics and microfluidic single-cell microscopy revealed unexpected, rapid protein and nucleic acid synthesis already in non-dividing cells. Additionally, the lag time to first division shortened as pulsing frequency increased. Here, we demonstrate that the first division from a non-dividing state occurs when the facilitating protein FtsZ reaches division-supporting concentration. A dynamic model quantitatively relates lag time to FtsZ synthesis from nutrient pulses and its protease-dependent degradation. Consistent with model predictions, lag time shortened when FtsZ synthesis was supplemented or protease inhibitors were added. Lag time prolonged when *ftsZ* was repressed or FtsZ degradation rate was increased. Thus, we provide a basis to quantitatively predict bacterial division using information about molecular determinants and the nutrient input.

## Main

The division of one cell into two daughters is a key feature of life, and we understand many molecular processes that achieve this fundamental biological event in different cell types. Less clear is the exact molecular basis to initiate the division process, especially in relation to nutrient input. Nutrition-related cues proposed as decision signals include protein^1^ or DNA^2^ concentrations, and metabolites that interact with the division machinery^3^. Current models of bacterial division focus on exponential growth conditions^4^ where nutrients are abundant. Typically, these models use phenomenological quantities such as biomass per cell as the decision input variable. For example, the adder model^5,6^ accurately predicts that bacteria will divide after a constant amount of biomass addition after birth for exponential growth.

Before bacterial cultures can divide exponentially, individual cells must first make the decision for the initial division from a non-dividing state, the typical situation for microbes in their natural environment^7^. Moreover, in many environments, non-dividing microbes receive nutrients only sporadically and in small quantities, such as in the gut^8^, soil^7^, ocean^9^, but often also in industrial fermentation processes^10^. The biomass per cell input variable is not sufficiently detailed to understand the decision process for the first cell division of a non-dividing state. Furthermore, the biosynthetic capabilities of starved cells are generally not well understood^11^. Hence, current models of cell division do not predict division timing for the widespread, naturally occurring sporadic nutrient conditions. Thus, open questions remain: How do cells decide the first division from a non-dividing state? Which molecular entities determine their decision?

Here, we studied the first division decision of starved *E. coli* under sporadic nutrient supply. We developed methodologies to measure division occurrence and metabolic activity of starved cells under sporadic pulsing. We found that cells rapidly synthesized proteins and nucleic acids from sporadic glucose. Additionally by quantifying division timing as a function of sporadic glucose pulse frequency, we deduced the FtsZ protein as the division determinant, built a quantitative model, and substantiated it with follow up experiments.

## Results

### The lag time to division shortens with glucose pulse frequency for a subpopulation

We developed three complementary systems (Fig. 1) – each with different advantages and providing robust cross-validation of each other – to controllably pulse nutrients to starved *E. coli* and measure division occurrence. Two of the systems (spin flask and plate reader) pulsed nutrients by dispensing a drop of defined volume at a programmed frequency to a starved culture. The drops were calibrated so that the final concentration, after the pulse mixed with the culture, was the same between the two systems. In the third system, bacteria attached to the bottom surface of a microfluidic chamber were suffused with flowed media and imaged over time, while a pressure system controller allowed a precise and rapid switch of flowing medium and similarly provided nutrient pulses to the bacteria.

**Figure 1:**
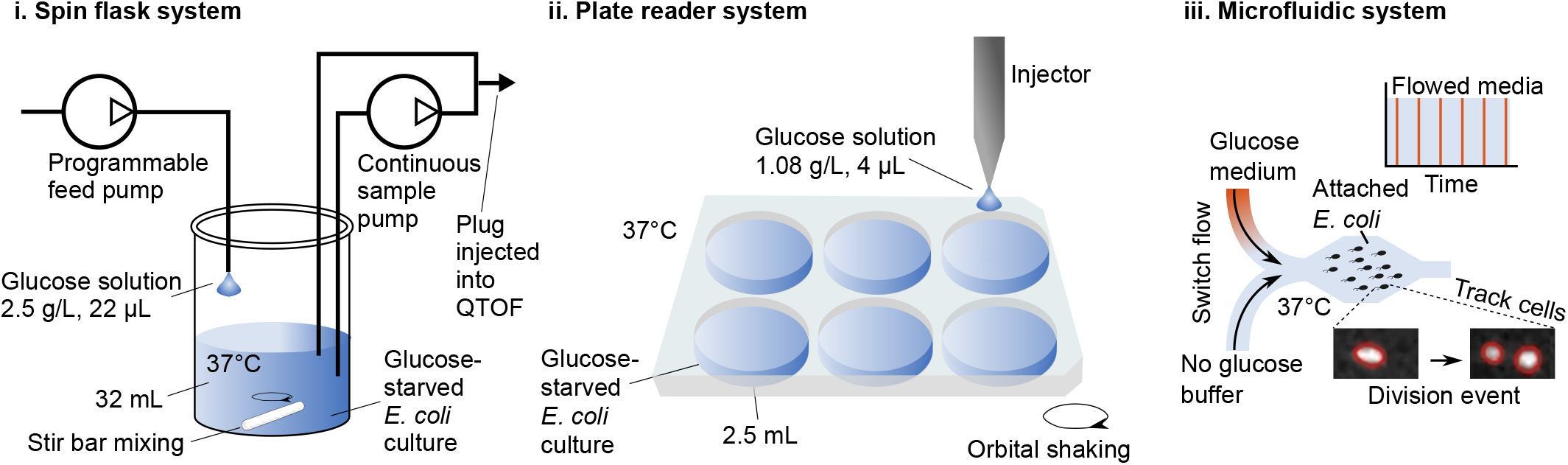
Schematics for nutrient pulse systems. Three separate systems were used to pulse glucose to starved *E. coli*. The spin flask system (i) and plate reader system (ii) provided glucose pulses at defined frequencies. In the real-time metabolomics^17^ configuration, another pump circulated culture and injected 2 μL of culture directly into a time of flight (QTOF) mass spectrometer every 10–15 s from the spin flask system. A microfluidic platform (iii) reproduced the pulse feeding and tracked division events. A pulsing period is defined as the time between the start of successive glucose medium exposures. During each pulse, glucose medium was flowed for 10 seconds, and the no glucose buffer was flowed in the intervening period.

How does sporadic nutrient availability empirically relate to the division decision? We focused on the case of limiting carbon and energy with sporadic glucose pulses. Glucose grown cells were starved for 2 h and then pulsed at controlled frequencies of 10 μM glucose with the spin flask and plate reader systems. Hereafter, we use the term time-integrated (TI) feedrate (abbreviated *f*, units: mmol glucose/g dry cell weight/hour) as the average rate of glucose fed over time normalized to the initial mass of cells in the culture. Our pulse frequency-modulated TI feedrates spanned the range from 0.1 mmol/g/h, which does not support division, to just above 1 mmol/g/h. All TI feedrates were well below the exponential growth consumption rate of *E. coli* (~10 mmol/g/h)^12^. The cultures were glucose limited throughout the experiments, as was verified by absent glucose accumulation after pulses (Supplementary Table 1). To assess division occurrence as a function of TI feedrate, we measured the optical density (OD). Strikingly, the transition (lag) time to cell division, i.e., from constant to increasing OD, was dependent on pulsing frequency (Fig. 2a and Supplementary Table 2) and not explained by the total glucose fed (Supplementary Figure 1). At TI feedrates below ~0.2 mmol/g/h (the critical rate), the OD did not increase within the first 6 hours of pulsing. Above ~0.2 mmol/g/h, OD increase was only observed after a TI feedrate-dependent lag time from the start of pulsing (Fig. 2a insets). For TI feedrates above ~1.0 mmol/g/h, OD increased immediately without a detectable delay.

**Figure 2:**
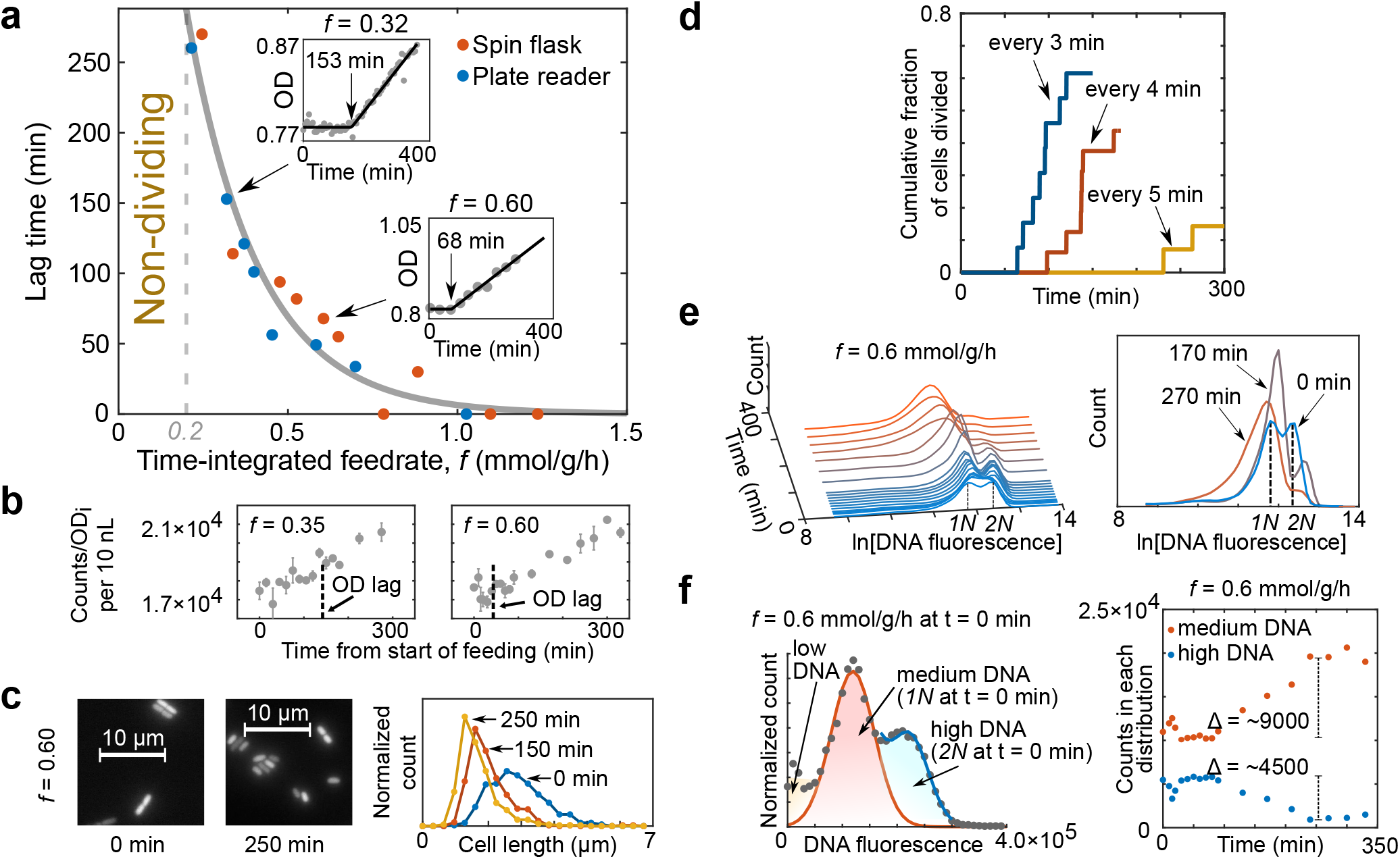
Lag time to division depends on the frequency of pulsed glucose for a subpopulation. **a**, After 2 hour starvation, *E. coli* cultures were pulse fed 10 μM glucose at varying frequencies using the spin flask and plate reader systems, and optical density (OD) was measured over time (**inset** example figures). Grey dots are OD measurements and the black lines are an empirical fit (see Methods). For separate experiments (*n* = 18), the lag time is plotted against the frequency, represented as the time-integrated (TI) feedrate *f* (mmol glucose/g dry cell weight/hour). An empirical fit (grey solid line, see Methods) was used to separate the lag (non-dividing) and dividing phases. All OD data is summarized in Supplementary Table 2. **b**, Normalized absolute cell counts versus time show linear increases after lag time for exemplary feedrates. Data are mean ± standard error of technical replicates (*n* = 2–3). Lag predicted from the empirical fit is indicated by vertical dotted lines. **c**, The average size of cells decreased after lag. Micrographs and cell length distributions (*n* > 400 per distribution) are shown for specific time points, with *f* = 0.6 mmol/g/h. **d**, Immobilized cells in the microfluidic experiment divided after a lag time that decreased with increasing glucose pulse frequency. The labeled times indicate the period, time between pulses, for a given experiment. **e**, Time course of the distribution of cellular DNA content. Sampled cells were stained with SYBR Green I and measured with flow cytometry over the course of a pulsing experiment (*f* = 0.6 mmol/g/h). Gating is shown in Supplementary Figure 2. The DNA content distribution over time is shown on the left side, and three specific time points are shown on the right. Within the first time point (t = 0), the highest distribution is taken to be high DNA content (2N), and the distribution at half of the 2N average was taken (1N) as medium DNA. **f**, DNA distributions were separated into medium (1N) and high DNA (2N). Distribution-specific estimated counts (see Methods) over time (*f* = 0.6 mmol/g/h) suggested that net division from high to medium DNA cells can explain the increase in cell counts and OD increase.

We confirmed that the OD increase reflects cell division by observing similar inflections in cell counts measured with flow cytometry occurring after lag times (Fig. 2b). Since the total glucose fed during the lag phase was calculated to be insufficient for the doubling of the biomass of all cells in the culture (Supplementary Table 2), we expected the average cell to become smaller. Indeed, microscopy demonstrated that the average cell size decreased after the onset of cell division (Fig. 2c). Lastly, we used the microfluidic platform to similarly pulse feed cells and visually track division events (Fig. 1a and Supplementary Video 1). Consistent with our previous observations, division started after a lag time that shortened with increasing pulsing frequency (Fig. 2d).

We noticed that not more than ~65% of the cells divided within 5 h in the microfluidic experiments, suggesting potential population heterogeneity. Furthermore, the initial linear increase in OD flattened before the initial OD was fully doubled (Supplementary Figure 3). Both observations suggested that primarily a subpopulation undergoes the division. We hypothesized that this subset of cells were further along in the cell cycle before the start of pulsing compared to the rest. Therefore, we resolved the cell cycle status during pulsing by flow cytometric analysis of the DNA content distribution (Fig. 2e). Before pulsing, two subpopulations existed, one with low (1N) and one with double DNA per cell (2N), as previously observed for *E. coli* in stationary phase^13^. Upon glucose pulsing, the 2N cells disappeared while the 1N population increased. The 2N population was likely in the D period of the cell cycle^7^, with sufficient DNA for division but limited nutritionally. Counting both 1N and 2N cells over time suggested that all division could be explained by 2N cells dividing into 1N (Fig. 2f).

### Pulsed glucose is used rapidly to synthesize biomass even without division

How is pulse-fed carbon utilized during the lag phase? In principle, it could be consumed by non-growth related maintenance requirements^14^ or stored for division. We defined maintenance as any consumed glucose not used directly for division, but rather for energetic costs such as protein turnover and sustaining cell integrity. We wondered whether maintenance was equivalent to and explained the critical rate (~0.2 mmol/g/h), meaning that only fed glucose exceeding the maintenance could be utilized for division. We, therefore, decomposed the TI feedrate, *f*, into division and maintenance terms by assuming a linear dependence of the division rate (*Ψ*, units: 1/h • [number of new and existing cells/number of existing cells]) on the TI feedrate^15^ (Fig. 3). The division rate was almost directly proportional to the TI feedrate, suggesting that the required maintenance (i.e. the y-intercept) is less than the critical rate (~0.2 mmol/g/h) and generally too small for precise measurement, as seen before in carbon-limited batch culture^16^. We conclude that most carbon pulsed during lag is stored for eventual division and that the critical rate is not explained solely by maintenance requirement.

**Figure 3:**
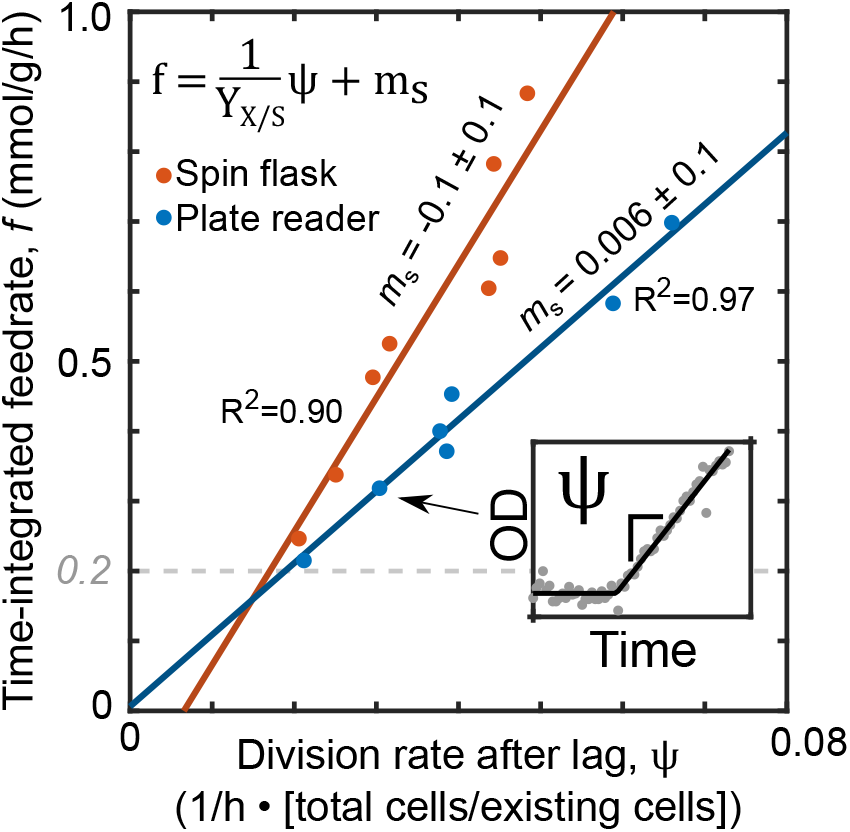
Maintenance metabolism alone cannot explain non-division. Linear decomposition of the TI feedrate (*f*) from data in Fig 2a separated the division (*Ψ*/*Y*_x/s_) and maintenance terms (*m*_s_):

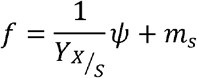 For the division term, the division rate (*Ψ*, units of 1/h • [number of new and existing cells]/[number of existing cells]) was calculated to be the slope after the lag ends (inset). For each pulsing system, the calculated yield (*Y*_x/s_, units of g DCW/mmol glucose • [number of new and existing cells]/[number of existing cells]) was constant and the extrapolated maintenance term (*m*_s_) was not significantly detected (*m*_s_ = −0.1 ± 0.1 mmol/g/h for spin flask system and *m*_s_ = 0.006 ± 0.1 mmol/g/h for the plate reader setup).

How do non-dividing cells process and potentially store sporadically pulsed carbon? To address this question we performed near real-time metabolomics at a resolution of 10–15 s during the glucose pulses^17^. A continuous sample pump circulated culture liquid and provided 2 μL of whole cells in medium to a flow injector with time-of-flight mass spectrometer. More than 100 different annotated metabolites were measured (Supplementary Table 1). We observed sharply defined pulse responses in the concentration of all detected central metabolic intermediates at TI feedrates of 0.06, 0.12, and 0.18 mmol/g/h (Fig. 4a and Supplementary Figure 4a) that did not support cell division (Fig. 2a). The concentration spike and the return to baseline levels within about 300 s strongly suggested that a wave of carbon rapidly flows through central metabolism. In sharp contrast, several building blocks of cellular biomass such as amino acids and nitrogen bases continuously increased between pulses and rapidly decreased immediately after each glucose pulse (Fig. 4a and Supplementary Figure 4b). Since these accumulated amino acids including phenylalanine cannot be degraded by *E. coli*, their depletion suggested a brief increase in protein synthesis with each pulse^18^ (Fig. 4b). The nitrogen bases, hypoxanthine and guanine, may be salvaged for new nucleic acid synthesis upon sudden access to carbon. These observations suggested that fed carbon rapidly sweeps through central metabolism into biosynthesis of amino acid and nucleotide monomers and leads to a period of increased protein and nucleic acid synthesis immediately after the glucose pulses, even in the absence of cell division. This observation echoed earlier work about net protein synthesis in lag phase before division^19^.

**Figure 4:**
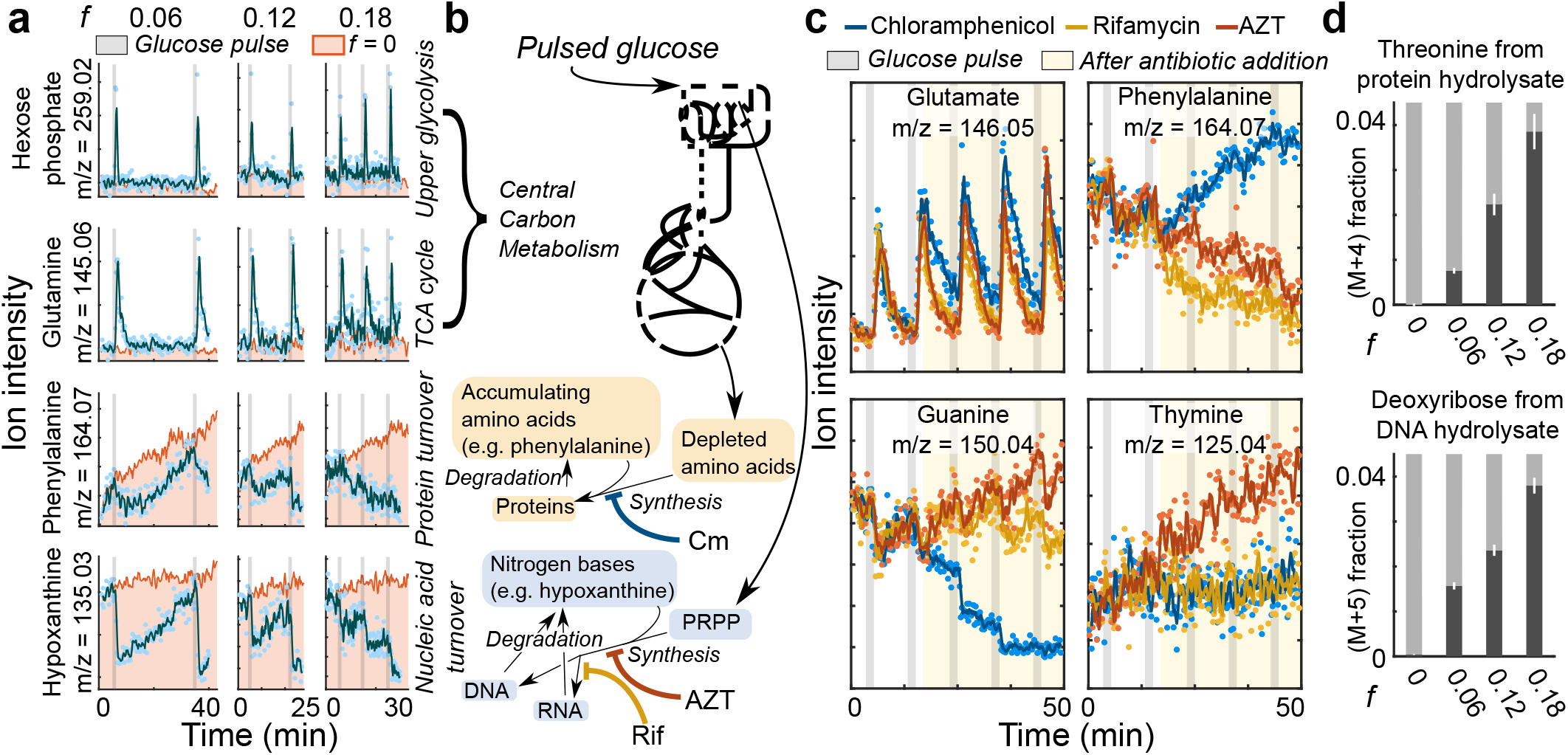
Non-dividing *E. coli* use pulse-fed carbon to make nucleic acids and protein. **a**, The spin flask system for glucose pulsing was connected to a real-time metabolomics platform. Traces of exemplary ions are shown that correspond to hexose phosphate, guanine, phenylalanine, and hypoxanthine for pulsing at non-division supporting frequencies of 0.06, 0.12, and 0.18 mmol/g/h. The TI feedrate is abbreviated as *f* (units: mmol glucose/g dry cell weight/hour). Glucose pulses are indicated by the grey bars, and the pink region shows a no pulse control. Dots are ion intensity measurements. Solid lines are a moving average filter of the measured ion intensity. For clarity, dots are not shown for *f* = 0 mmol/g/h condition. **b**, A metabolic scheme describing the propagation of fed glucose. Pulsed glucose is hypothesized to pass through central carbon metabolism and then be converted to downstream pathways including amino acid synthesis and nucleic acid synthesis. For nucleic acid synthesis, glucose is converted to the intermediate PRPP, which then can combine with nitrogen bases to form nucleotides for nucleic acid synthesis. Different pathways can be blocked with antibiotics. Color scheme used here accords to Fig. 4c. **c**, Influence of antibiotics that inhibit macromolecular synthesis at the non-division TI feedrate of 0.18 mmol/g/h. Antibiotics were added one minute after the second pulse (yellow region). Four different ions are shown corresponding to glutamate, phenylalanine, guanine, and thymine. Chloramphenicol (blue) inhibits protein biosynthesis, rifamycin (orange) inhibits RNA polymerase, and azidothymidine (AZT; red) inhibits DNA synthesis. Ion traces with negative control (*f* = 0.18 mmol/g/h, no antibiotics) are shown in Supplementary Figure 5. **d**, Percentage of labeled threonine and deoxyribose from protein and DNA hydrolysate shows *de novo* protein and DNA synthesis in non-dividing cells. After 6 hours of pulsing uniformly labeled ^13^C-glucose, cultures were lysed, and their macromolecules were washed free of latent metabolites and hydrolyzed to monomers. Labeling data are presented as the mean ± standard error of independent biological replicates (*n* = 3, all pairwise *P* < 0.02 as determined by one sided Student’s t-test). All ion data is available in Supplementary Table 1, and labeling data of all measured amino acids is available in Supplementary Table 3.

To confirm protein and nucleic acid synthesis from fed carbon in non-dividing cells, we repeated the glucose pulsing and blocked macromolecule synthesis by adding antibiotics one minute after the second pulse to curtail carbon to specific metabolic sectors (Fig. 4bc and Supplementary Figure 4c). Upon addition of the ribosomal inhibitor chloramphenicol, the depletion of five measured amino acids including glutamate and phenylalanine was slowed compared to addition of other antibiotics. Conversely, guanine but not the amino acids exhibited a similar effect upon challenge with rifamycin and azidothymidine, which limit RNA and DNA synthesis, respectively^20^. The DNA-specific nitrogen base thymine, as expected, accumulated only upon azidothymidine addition. To directly demonstrate incorporation of fed glucose into biomass macromolecules at non-division frequencies, we performed pulse experiments with uniformly labeled ^13^C-glucose for 6 h. Increasing fractions of labeled threonine (M+4) and other amino acids in extracted and hydrolyzed protein confirmed *de novo* protein synthesis (Fig. 4d and Supplementary Table 3). Likewise, increasing labeled fractions of deoxyribose (M+5) from hydrolyzed DNA substantiated the use of pulsed carbon for *de novo* DNA synthesis (Fig. 4d) through the PRPP intermediate as shown previously^17^. Although glycogen is a storage form of glucose^21^, much less labeling was found in glycogen hydrolysate (Supplementary Table 3). Lastly, we tested whether macromolecular synthesis occurred primarily in the 2N population using single cell microscopy under microfluidics with nutrient pulsing (Supplementary Figure 6). We separated populations of cells into dividing (all 2N) and non-dividing. Dividing cells synthesized more biomass and protein before division compared to non-dividing cells. Specifically, the cell elongation and GFP synthesis rates were higher in dividing cells (dividing cell extension rate of 0.0086 ± 0.0014 μm/min, dividing GFP synthesis rate of 9.2×10^−5^ ± 1.8×10^−5^ Norm. GFP/min versus non-dividing cell extension rate of 0.0033 ± 0.0013 μm/min, non-dividing GFP synthesis rate of 6.7 ×10^−5^ ± 1.9×10^−5^ Norm. GFP/min). Collectively, antibiotic challenges, ^13^C-labeling, and microfluidics support our hypothesis that fed carbon is assimilated into protein and nucleic acids in non-dividing cells during the lag phase.

### FtsZ synthesis and degradation determines the division occurrence

Next, we asked what determines division occurrence. We conjectured that a defined stoichiometry of key macromolecules (e.g. division proteins, DNA) commences the division event. Since pulse-fed glucose is converted into protein, RNA, and DNA in non-dividing cells, we posit that a specific macromolecule may stoichiometrically limit the division. Given that the lag time to the first division is a function of the pulse frequency, the most parsimonious explanation is that the limiting macromolecule(s) are synthesized after the pulse for a brief period and constitutively degraded (Fig. 5). This means that longer time between pulses results in more degradation and greater total glucose requirement to reach division, which is consistent with our data (Supplementary Figure 1). The competing synthesis and degradation also can explain the critical rate (~0.2 mmol/g/h); a critical rate would exist where the synthesis and degradation rates of the limiting entity are equal (*f*_3_ in Fig. 5). Since proteins are the most abundant macromolecules^22^ and because their degradation kinetics are consistent with the time scales observed^23^, we hypothesized that the limiting, determining entity is a degraded protein. A key aspect of this theory is amenable to experimental validation: the lag time should be reduced by abrogating protein degradation with chemical protease inhibitors. Specifically, we added a cocktail of protease inhibitors at the onset of pulse feeding, using *f* = 0.28 mmol/g/h for which the usual lag time was about 200 min. Consistent with our hypothesis of continuous degradation of one or more proteins that limit division, treatment with protease inhibitors reduced the lag time by 30% (Fig. 6a).

**Figure 5:**
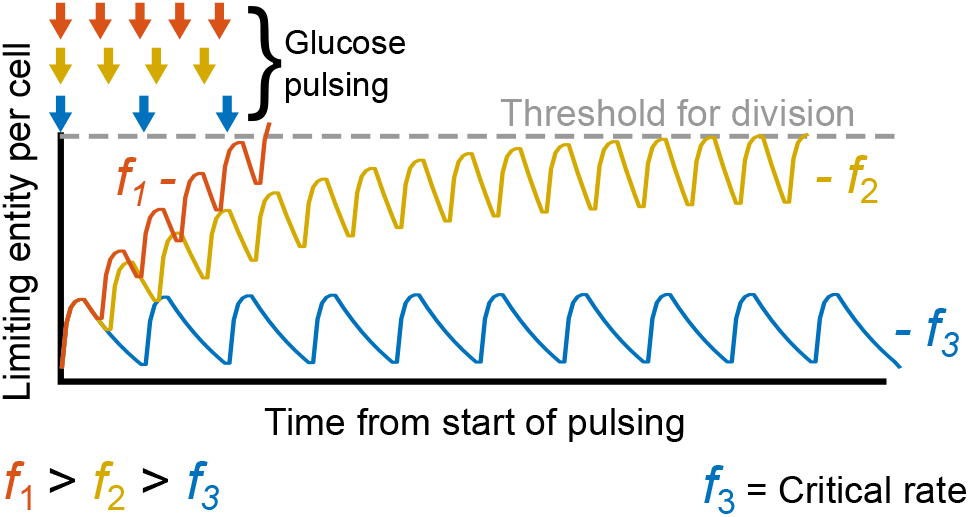
The limiting, degrading entity hypothesis. The dependence of lag time on glucose pulse frequency can be explained with constitutive degradation of the limiting entity. In the model shown, the entity abundance is synthesized with each glucose pulse and depletes constitutively. Three example frequencies (*f*_1_ > *f*_2_ > *f*_3_) are shown where slight changes in period time dramatically changes the time for the entity to reach the threshold needed to engender division. When synthesis and degradation of the entity are equal, the TI feedrate is at the critical rate (*f*_3_). Arrows indicate the glucose pulse frequency.

**Figure 6:**
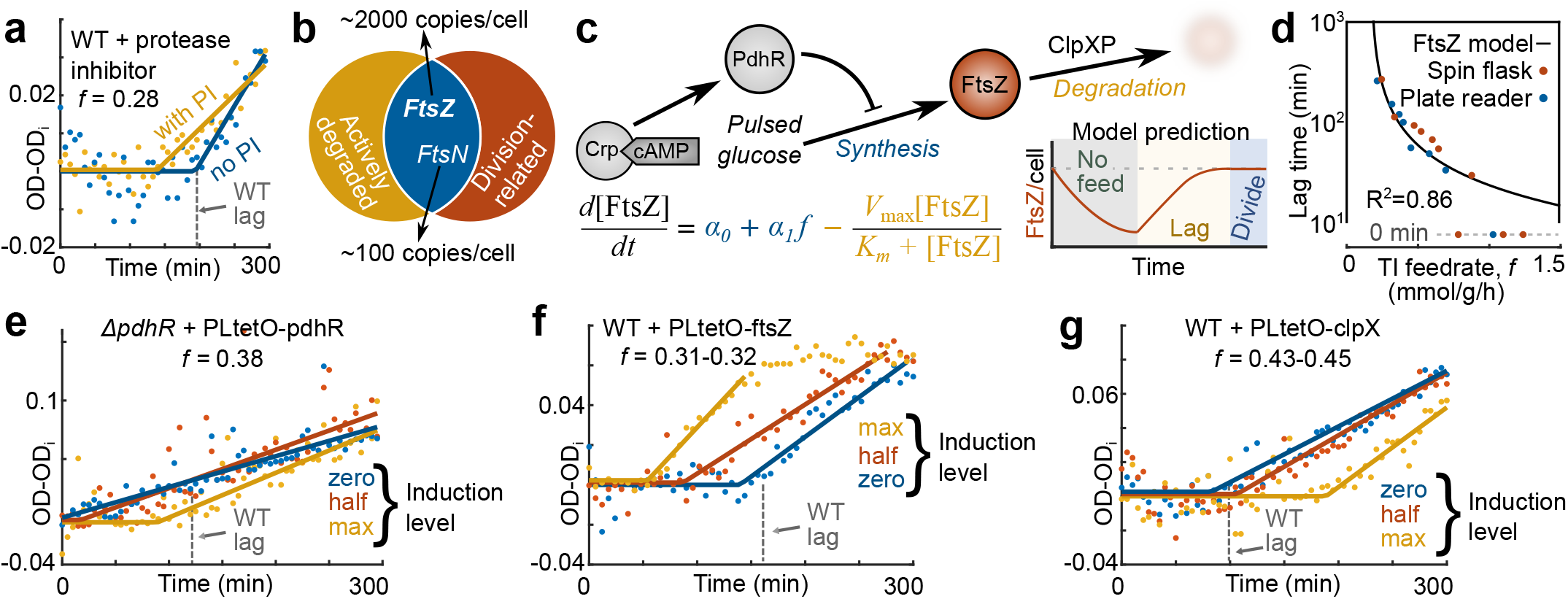
Dynamics of FtsZ abundance determines the division timing. **a**, Pulsing experiment was repeated in the presence of protease inhibitor (PI) that reduced the lag time for a given TI feedrate (*f* = 0.28 mmol/g/h). The TI feedrate is abbreviated as *f* (units: mmol glucose/g dry cell weight/hour). Wild-type lag (from Fig. 2a empirical fit) is indicated by the dotted grey line. **b**, The sets of proteins that are actively degraded and division-related intersect at FtsZ and FtsN. **c**, A schematic of how FtsZ abundance changes. FtsZ is repressed by the transcriptional factor, PdhR. PdhR is activated by Crp-cAMP. FtsZ is also degraded primarily by the ClpXP protease complex. An approximate FtsZ threshold model poses a basal synthesis rate (α_0_), a feedrate-dependent synthesis (α_1_*f*), and a degradation term (the Michaelis Menten term) to explain changes in FtsZ abundance with and without pulsing. Per the model, FtsZ would deplete via degradation during starvation, be synthesized with glucose pulsing, and engender division when its abundance reaches the threshold concentration. **d**, Analytical solution of the model (Supplementary Information) plotted against data from Fig. 2a (R^2^ = 0.86). Lag time axis is log scaled. **e**, Genetically induced titration of PdhR in a *pdhR* mutant reintroduced lag commensurate with expression level for a given TI feedrate (*f* = 0.38 mmol/g/h). Induction level corresponds to the amount of doxycycline (max – 50 ng/uL, half – 10 ng/uL, and zero 0 ng/uL) added at the onset of starvation. **f**, Lag time reduced with synthesis levels of titrated FtsZ in the wild-type strain at *f =* 0.31–0.32 mmol/g/h. **g**, Lag time increased with titrated synthesis of ClpX in wild-type cells at *f* = 0.43–0.45 mmol/g/h.

To identify the putative division limiting protein for division, we considered the known set of degraded proteins in *E. coli*^24,25^, approximately 7% of the proteome. When we intersected the degrading protein set to the set of proteins involved in cell division^26^, only FtsN and FtsZ remained (Fig. 6b). Given that FtsN has very low abundance of around 100 copies per cell^27^, we focused on FtsZ. FtsZ forms the division ring that septates a mother cell into two daughters^28^. FtsZ is transcriptionally repressed by PdhR^29^, which is activated by the global transcriptional regulator Crp-cAMP^30^ (Fig. 6c). Since Crp-cAMP regulation is highly active during carbon starvation in *E. coli*^31^, one would expect *ftsZ* to be repressed during starvation and in the lag phase. Indeed, genetic disruption of *ftsZ* repression by deleting *crp* or *pdhR* entirely abrogated the non-division phase, as cells divided without lag upon pulsing (Supplementary Figure 7). These results suggest that FtsZ limits division and is synthesized during each pulse while being continuously degraded until its concentration reaches a level that supports division (Fig. 6c inset). We tested the plausibility of this hypothesis by developing an approximate, smoothed dynamic model:

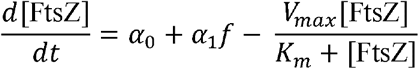

The model accounts for the basal synthesis (α_0_), pulsing-dependent synthesis (α_1_*f*), and degradation (the Michaelis-Menten term) of FtsZ. We parameterized the model based on literature values mostly specific to FtsZ, the strain, and the media used^23,27,32^ (Supplementary Information). Despite fitting just a single parameter α_1_, the model reproduced non-zero lag times remarkably well (R^2^ = 0.86), supporting the role of FtsZ as the limiting entity for division (Fig. 6d).

Our model postulates that FtsZ abundance depletes monotonically during starvation and increases upon glucose pulsing. Since resolving FtsZ abundance changes within a single pulse interval requires intractable sensitivity (FtsZ abundance changes ~1% between pulses, Supplementary Information), we monitored FtsZ abundance changes over longer periods with immunoblotting. Pulsing for 16 h at non-division inducing TI feedrates yields several-fold higher FtsZ concentrations compared to 16 h starvation (Supplementary Figure 8), confirming that FtsZ is indeed one of the proteins synthesized from the glucose pulses under starvation. Genetic deletion of *clpX* or *clpP* similarly increased FtsZ concentrations even under full starvation, confirming *in vivo* that the ClpXP protease complex degrades FtsZ during glucose starvation. This result is consistent with earlier *in vitro* evidence for ClpXP-based degradation of FtsZ^32^.

Observed synthesis and degradation of FtsZ alone, however, cannot establish division determinacy because many proteins are likely synthesized with glucose pulses and degraded. Instead, the model proffered clear, falsifying experiments to test FtsZ’s candidacy as the limiting entity. We first titrated FtsZ synthesis, in effect modulating specific parameters while holding initial/division conditions, TI feedrate, and other parameters. At a TI feedrate of 0.38 mmol/g/h, a mutant strain with *pdhR* deletion that lacked FtsZ transcriptional repression divided without a lag phase, but the lag phase was gradually restored upon plasmid-based expression of PdhR (decreasing α_0_ and α_1_) (Fig. 6e). Similarly, direct plasmid-based supplementation of FtsZ (increasing α_0_ and α_1_) in the wild-type reduced the lag time with increasing induction levels for a given TI feedrate (Fig. 6f). The causal role of protein degradation was tested by modulating the FtsZ degradation rate through plasmid-based overexpression of ClpX. ClpX abundance is known to be rate-limiting for ClpXP-based degradation^33^; therefore, supplemented ClpX should increase the FtsZ degradation rate (increasing *V*_max_). Consistent with our hypothesis, lag times prolonged at a given TI feedrate with increasing ClpX expression in *E. coli* (Fig. 6g). We conclude that all titration experiments affecting the synthesis and degradation parameters are consistent with FtsZ division determinacy.

To exclude the possibility that also other division proteins are limiting, we titrated FtsA, FtsB, FtsL, and FtsN (Supplementary Figure 9). Overexpression of the former three did not affect the lag time, but at the highest induction level FtsB and FtsL increased the division rate once the lag time ended. FtsN overexpression exhibited a more complex phenotype. While the highest induction level appeared to reduce the lag time, it also had a deleterious effect resulting in only a small increase in OD, and thus presumably division of only very few cells. FtsN cannot, therefore, be the sole limiting factor. The reduced lag with supplemented FtsN may be explained by a population minority where FtsZ abundance is enough and FtsN is limiting. Alternatively, supplemental FtsN could affect FtsZ polymerization, degradation, or the separation of cell^34^, thus affect lag through the FtsZ determinacy model. We conclude that the negative controls do not falsify the FtsZ determinacy model, but other division proteins may influence division through their interaction with FtsZ^35^ or may potentially be limiting in a smaller fraction of cells where FtsZ is abundant enough.

Finally, we wondered whether FtsZ-limited division is specific to pulsed glucose or a more general mechanism that links the nutritional status to the first cell division. For this purpose, we tested the influence of FtsZ overexpression on the lag phase upon pulsing carbon-starved *E. coli* with the gluconeogenic carbon sources glycerol and acetate and nitrogen-starved cells with the nitrogen source ammonium (Supplementary Figure 10). In all cases, FtsZ overexpression reduced the lag phase akin to the glucose case. Thus, our results suggest that the balance between FtsZ synthesis and protease-mediated degradation is a general control mechanism for the first cell division during sporadic nutrient availability for a variety of different nutrients.

## Discussion

The rapid, untrammeled biomass synthesis in non-dividing, starved cells surprised us. Starved cells are expected to throttle metabolism and *de novo* biosynthesis (transcription, translation, and DNA replication) due to the stringent response effects^11^ and, therefore, cease accumulating biomass^36^. Our expectation and the implicit one from earlier work anticipated that the metabolite pools must first replenish to continue the cell cycle and biosynthesis. Our measurements demonstrate that over a period of a few hours, glucose-starved *E. coli* maintain a high anabolic and catabolic capacity. Furthermore, measured central carbon metabolite pools recrudesce and deplete within seconds, meaning even minuscule glucose passes through quickly. The limitation for division occurred on the protein level and not a specific metabolite, echoing recent work that argues the protein production and not metabolic activity limits cell cycle progression^37,38^.

Additionally, we demonstrated that, under conditions of sporadically available nutrients, the dynamics of FtsZ concentration primarily determines the timing of first cell division in *E. coli*. The hypothesis that FtsZ concentration is a determinant of division has been proposed before^1^, but was rejected in experiments conducted under exponential growth conditions^39,40^, where glucose consumption is saturated at a high consumption rate of 10 mmol/g/h^12^. We further demonstrated with a computational model that a FtsZ-driven mechanism can quantitatively explain the timing of division for low levels of nutrient consumption (below ~1 mmol/g/h). This further suggests that the determinant of division in *E. coli* is consumption rate dependent. Additionally, to the best of our knowledge, only phenomenological models such as the adder model^5,6^ have to date been proposed to quantitatively predict the timing of division. Here we have formulated the underpinnings of a mechanistic, biochemical model that provides quantitatively deeper understanding of the first decision to divide in relation to nutritional input.

## Methods

No statistical methods were used to predetermine sample size. The experiments were not randomized. The investigators were blinded to some sample measurements and outcome assessment.

### Strains and plasmids

*E. coli* BW 25113 from the Keio collection^41^ was used as the wild-type (WT) strain. Kanamycin markers were excised from the Keio knockout strains *crp*, *pdhR* using pCP20 and verified using PCR^42^. All strains are listed in Supplementary Table 4 and available from authors on request.

Plasmids are listed in Supplementary Table 5, and all GenBank files are available in Supplementary Data 1. All plasmids originating from this study were designed using j5 software^43^, assembled using Gibson-based techniques^44^, and sequence verified (Microsynth). Briefly, the titratable pJKR-L-tetR plasmid^45^ was used as a template where the sfGFP sequence was replaced with *pdhR*, *ftsZ*, *clpX, ftsA, ftsB, ftsL*, and *ftsN*. The plasmid pJKR-L-tetR was a gift from George Church (Addgene plasmid # 62561). For microfluidic experiments, the plasmid epd-icd^46^ was used for constitutive GFP expression. All plasmids originating from this study are available from AddGene (Article No. 25280).

### Cultivation, pulse feeding, and chemical concentrations

Cultivation procedure was followed as in an antecedent study^17^. All cultivation was performed at 37°C in shaker unless stated otherwise. Briefly, the day before pulsing, cells from freezer stock were cultivated in LB media for 3-5 h, then diluted 1:20 into 5 mL total of M9 media (see ^17^ for recipe), and cultivated for 4-5 h to OD 0.1-0.2. 500 uL of the 5 mL inoculum was dispensed into 35 mL of M9 media and cultivated overnight at 30 °C. The next day, cultures were typically at OD 0.2-0.3 and were then moved to 37°C and cultivated until OD 0.8-1.2. At this point, cells were pelleted by centrifugation (3 minutes at 5000 rpm) and resuspended in 32 mL of diluted M9 media without glucose (1:8 dilution with filtered water). This point signified the start of starvation. Cultures were then cultivated without glucose for 2 h before the start of pulsing.

Glucose pulsing was accomplished using two systems (Fig. 1a). With the spin flask system, an IDEX Corporation Ismatec MCP 404 pump was programmed to dispense 22 μL of 2.5 g/L glucose solution to a 32 mL culture within a Schott bottle. Starved cultures were transferred to Schott bottle just before the start of pulsing. Frequency/flow rate was controlled by setting pause time between dispensations. Cultures were constantly mixed using a stir bar and maintained at 37 °C by submergence into a water bath. Optical density (OD) was measured in a Pharmacia Novaspec II spectrophotometer. In the plate reader system, a Tecan Reader Infinite 200 with injector was programmed to dispense 4 μL of 1.08 g/L glucose solution to a 2.5 mL culture in 6 well plates ((Thermo Fisher Scientific). Plate reader cultivations were performed at 37 °C and with orbital shaking at maximum amplitude. An empirical function was used to convert OD measurements from the plate reader system to the spectrometer one.

Final concentrations of antibiotics were as follows: 100 μg/mL of ampicillin, 34 μg/mL of chloramphenicol, 50 μg/mL of rifamycin, and 100 ng/mL of azidothymidine. For plasmid titration experiments, doxycycline was added to the media at the onset of starvation. Each inducer concentration was cultivated in separate shake flasks during starvation. A titration curve is shown in Supplementary Figure 11 for the plasmid expressing GFP. 50 ng/mL working concentration of doxycycline was used for maximal synthesis, 10 ng/mL for half synthesis, and none for zero synthesis. For the protease inhibitor experiment, a cOmplete EDTA-free Protease Inhibitor Cocktail (Roche) tablet was dissolved in 2 mL of diluted media to form a stock solution. The stock solution was diluted 1:10 in M9 media without carbon source, inoculated with wild-type *E. coli*, and cultivated for two days at 30 °C to catabolize latent carbon within the cocktail solution. Cultivation was then pelleted, and the supernatant was collected and sterile filtered. Filtered, spent protease inhibitor solution was kept at 4 °C for no more than one day before experiment. Spent protease inhibitor solution was warmed to room temperature and added 1:10 (total dilution of 1:100 from stock) at the onset of pulse feeding for the experimental condition. For the negative control condition, spent diluted M9 was added instead.

### Flow cytometry and DNA distribution analysis

Flow cytometry procedure was extended from a previous study^47^. Two to three 5 μL samples were taken at every time point and diluted 1:10 in stain solution (filtered, spent media with 1:10000 SYBR Green I and 1:5 propidium iodide). Stained samples were incubated for 10 to 15 minutes, diluted 1:100 in filtered, spent media (total dilution of 1:1000), and then immediately measured in a BD Accuri C6 analyzer (BD Biosciences). 10 uL of diluted sample were injected at each time point and the first three time points were used to calibrate the expected number of events (*E*_i_). Absolute counts for each sample (*C*_s_) were calculated by accounting for clogging in the sample injection port using the equation *C*_s_ = *E*_cells,s_/*E*_total,s_. *E*_i_ where *E*_cells,s_ is the events in the gate (shown in Supplementary Figure 2) for a given sample, and *E*_total,s_ is the total number of events in a sample. The instrument settings were the following: Flow rate: slow; Threshold limits: 800 on SSC-H, 300 on FL1-H. All data was exported to CSV tables, and then gated and analyzed in MATLAB 2015b (Mathworks). FL1-H was used for DNA fluorescence. DNA distribution peaks were separated by fitting a combination of two normal distributions in MATLAB 2015b.

### Fluorescence microscopy and image analysis

5 μL samples were taken at every time point and diluted 1:10 in stain solution (filtered, spent media with 1:10000 SYBR Green I). After 10 minutes, 12 μL of stained sample was deposited onto 2-mm-thick layer of 1% agar on top of a microscope slide. The agar with samples were dried under air flow, and a cover slip was placed and glued. The cells were then immediately imaged (phase and fluorescence) using a Nikon Eclipse Ti inverted epifluorescence microscope equipped with a CoolLED PrecisExcite light source and a Nikon 100× oil immersion objective. Filters used for fluorescence imaging of SYBR Green I were 505 nm (excitation) and 545 nm (emission). The exposure time was set to 12 ms. Cell lengths were calculated using the Straight and Segmented line tools in ImageJ. At least 400 cells were measured for each time point.

### Microfluidics and analysis

The WT strain with epd-icd^46^ (constitutive GFP expression) was used for all microfluidic experiments. Non-glucose buffer was diluted M9 media conditioned for 2 hours with starved cells and then filtered. Glucose media was the non-glucose buffer supplemented with 200 μM glucose. Cells were exposed to non-glucose buffer for at least 2 hours before glucose exposures to provide initial starvation. Microfluidic channels were 100 microns wide (where the cells were imaged) and 60 microns deep, with two inlet ports, a 5-pointed junction, and two outlet ports. A pressure control system (Fluigent) allowed control of the duration and frequency of the glucose pulses. Before injecting the cells, the microfluidic devices were incubated with poly-L-lysine (Sigma, P8920; concentration 0.01% w/v) for 15 minutes to enhance the attachment of bacteria to the bottom glass surface of the channels^48^. All experiments were performed using a Nikon Ti-E inverted epifluorescence microscope equipped with Andor Zyla sCMOS camera, LED light sources (wavelengths 395, 440, 470, 508, 555, and 640 nm), a CAGE (LIS) incubator to maintain temperature at 37 °C, and a Perfect Focus System to reduce focal drift during long acquisition times. Image analysis was performed in MATLAB (Mathworks) using in-house cell tracking and identification algorithms. For calculations of GFP synthesis rate and cell extension rate, linear fitting was used on the data points for each cell before division.

### Real-time metabolomics profiling, annotation of ions, and data normalization

Whole cell broth, real-time metabolic profiling procedures were followed as in ^17^. The ion annotation method is described in ^49^. Ion suppression effects stemming from antibiotic addition are adjusted for as described in ^17^. All ion intensity data were Z-normalized and aligned (set *Z*) using the formula:

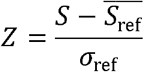

*S* is the raw ion counts, 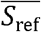 is the average of the reference set, and *σ_ref_* is the standard deviation of the reference set. For comparison to the non-pulsing condition (*f* = 0 mmol/g/h), the first five minutes were used for the reference set. For the antibiotic perturbations, the first ten minutes were used for the reference set. All annotated ion data before Z-normalization are available in Supplementary Data 2.

### Generation of washed lysates

Using the spin flask system, natively labeled cells were fed at TI feedrates of 0, 0.06, 0.12, or 0.18 mmol/g/h for 6 hours with uniformly-labeled ^13^C glucose. After the 6 hours, 25 mL of the culture were sampled, pelleted, and supernatants were discarded. Pellets were stored at −80°C until extraction. Cells were lysed via resuspension into 2.5 mL of B-PER solution (Thermo Fisher Scientific) and room temperature incubation for 10 minutes. 750 uL of lysates were clarified and spun through 10 kDa size exclusion columns (Merck Millipore Ltd.). The flow through was discarded, and the retentate was resuspended in 200 μL of filtered ddH_2_O. In total, three such spin-wash steps were performed and the final, washed retentate was used for further measurement.

### Protein hydrolysis and measurement

For protein hydrolysis and measurement, a previous protocol^50^ was extended. The washed lysate was adjusted to 6 N by HCl addition. Acidified lysates were incubated for 1 hour at 110° C and then dried under airflow at 65 °C. Dried samples were silylated by dissolution in 50 μL dimethylformamide and then added to 100 μL L N-tert-butyldimethylsilyl-N-methyltrifluoroacetamide with 1% tertbuthyldimethylchlorosilane. Reactions were then incubated at 85 °C for 1 hour. Products were measured on a 6890 GC combined with a 5973 Inert SL MS system (Agilent Technologies). Labeled fractions were adjusted for native isotope abundance^50^.

### DNA hydrolysis and measurement

For measurement of deoxyribose derived from purified DNA, the PureLink Genomic DNA Mini (Thermo Fisher Scientific) kit was used to isolate DNA from the cleaned lysate. 0.5 to 1.0 μg of DNA was then hydrolyzed to nucleosides using the EpiQuik One-Step DNA Hydrolysis Kit (Epigentek Group Inc). Reaction products were diluted to 100 μL with filtered water and spun through a size exclusion column. The flow through was directly measured on a 5500 QTRAP triple-quadrupole mass spectrometer in positive mode with MRM scan type (AB Sciex). Nucleoside standards were used for compound optimization. The deoxyribose-containing fragment of deoxyadenosine was measured for labeling fraction using SIM (*m*/*z* 252.3 > 117.2 through 257.3 > 122.2).

### Glycogen hydrolysis and measurement

For measuring glucose originating from glycogen, a previous method^51^ was extended. Washed lysate was acidified to 1 N by HCl addition in 300 μL total volume and incubated at 110° C for 1 h to hydrolyze polysaccharides. Samples were cooled on ice and neutralized with 84 μL of 3 N NaOH and then separated in size exclusion filters (10 kDa). The flow through was collected and dried in a SpeedVac setup (Christ) and precipitated with 500 μL cold ethanol. The ethanol resuspension was pelleted, and then the supernatant was separated and dried overnight in the SpeedVac. Samples were dissolved in 50 μL of pyridine with 2% hydroxylamine hydrochloride and incubated for 1 h at 90 °C. Samples were cooled to room temperature, and 100 μL of propionic anhydride was added. Mixed samples were incubated for 30 minutes at 60 °C and then measured on the aforementioned GC-MS system.

### Immunoblotting

For sampling, 500 μL of culture was collected, pelleted, and the supernatant was decanted. Samples were then immediately frozen at −20 °C for no more than one week before blotting. On day of blotting, samples were resuspended in 50 μL B-PER solution (Thermo Fisher Scientific) and incubated with shaking at room temperature for 10 min. Samples were pelleted and protein concentration was determined by Bradford assay (Biorad) according to supplied protocol. 1.5 μg total protein was loaded into each well of a 4-12% polyacrylamide gel (Sigma), electrophoretically separated, and transferred to a nitrocellulose membrane (GE Healthcare). Membranes were blocked using TBS-T buffer with 5% nonfat dry milk (Coop) for 1 h. Then membranes were consequently incubated with prokaryotic Anti-FtsZ primary antibody (Agrisera, Product No. AS10715) at 1:2000 dilution in TBS-T with milk overnight (4 °C with agitation). The membrane was then washed three times with TBS-T and incubated with HRP-conjugated anti-rabbit secondary antibody (Millipore, Product No. AP307P) at 1:10000 dilution in TBS-T with milk. Secondary incubation was conducted for 1 h, and then the membrane was washed with TBS-T three times. The membrane was then embrocated in Amersham ELC Prime Western Blotting Detection Reagent (GE Healthcare) to product specifications. After 5 min, the membrane was imaged first under bright-field to visualize ladder lanes and then with chemiluminescence measurement for protein bands. All imaging was done with a gel imaging station (Bucher Biotec). Ladder lanes were appended to the image with protein bands using GIMP software with exact pixel alignment. Band quantification was performed with MATLAB R2015B (Mathworks).

### Calculations and fitting

MATLAB R2015B (Mathworks) or Python 2.7 were used for all calculations, fitting (using the *fitnlm* function in MATLAB), and data analysis. Optical density was converted to gram dry cell weight (DCW) with the conversion 1 OD = 0.4 g DCW/L, as determined for the strain and spectrophotometer specifically^52^. For lag time (*t*_lag_), growth rate (*μ*), and initial cell amount (OD_i_) calculations, a threshold linear fit was applied to each OD versus time (*t*) plot:

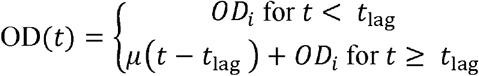

OD values 3 standard deviations above the mean of each dataset were excluded from fits.

To empirically separate the non-dividing and dividing phases (Fig. 2a), a threshold exponential decay fit was used:

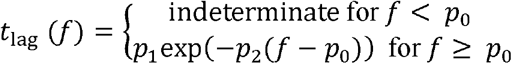

Correlation of coefficient (R^2^) for the FtsZ model was calculated after log_10_ transformation of the lag times. Data that had 0 minute lag time were excluded from the analysis.

### Code availability

All code used for calculations and figure generation is available without restriction in Supplementary Data 2 or at https://github.com/karsekar/pulsefeeding-analysis.

### Data availability

All data used in figures except from flow cytometry are available in the Supplementary Data 2 or at https://github.com/karsekar/pulsefeeding-analysis. Flow cytometry data is available at https://doi.org/10.5281/zenodo.1035825. Any other data is available from the authors on reasonable request.

## Acknowledgements

We thank the Sauer laboratory members, B. Towbin, W. Bothfeld, H. de Jong, M. Christen, R. Naisbit, E. Secchi, and E. Slack for helpful conversations, technical assistance, and manuscript feedback. We thank the ETH Flow Cytometry Core Facility for access, advice, and training for flow analyzer measurements.

## Author contributions

U.S. and T.F. conceived the study. K.S. and U.S. designed the experiments. U.S., R.S., and M.B. supervised the work. K.S. developed and performed the spin flask, plate reader, flow cytometry, microscopy, real-time metabolomics, immunoblotting, and molecular cloning procedures. K.S., T.F., and M.F.B. developed and performed the labeling experiments. J.N. and V.I.F. developed the microfluidic platform. R.R. performed the microfluidic experiments. K.S., R.R., E.N., and T.F. performed the calculations and analyzed the data. K.S. and E.N. developed the FtsZ model. K.S., U.S., E.N., and R.R. wrote the manuscript. All authors reviewed and approved the manuscript.

## Author information

Reprints and permissions information is available at www.nature.com/reprints. The authors declare no competing financial interests. Readers are welcome to comment on the online version of the paper. Correspondence and request for materials should be addressed to U.S.

**Supplementary information** is available in the online version of the paper.

---

**Supplementary Figure 1:**
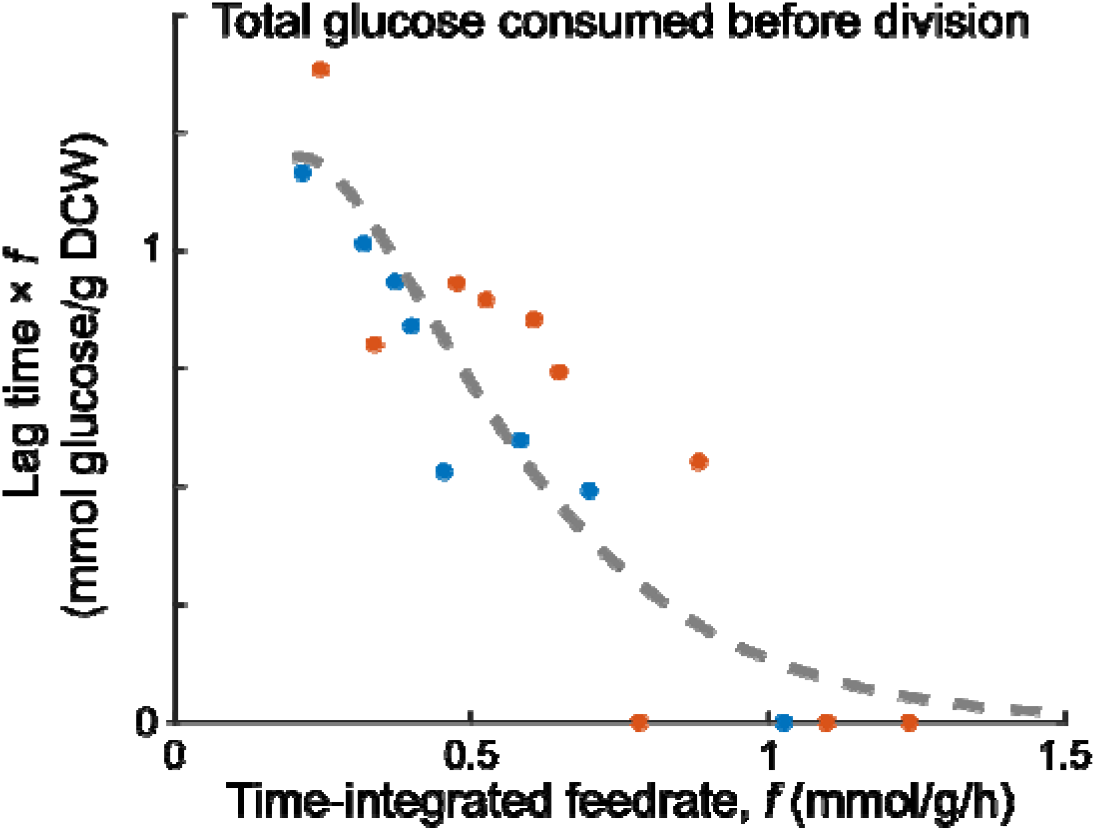
Total glucose fed during lag does not explain division occurrence. Data from Fig. 2a was replotted to total amount of glucose fed before division (lag duration times the TI feedrate) versus the TI feedrate. The total amount of glucose needed to trigger division is not constant and increases for decreasing TI feedrate.

**Supplementary Figure 2:**
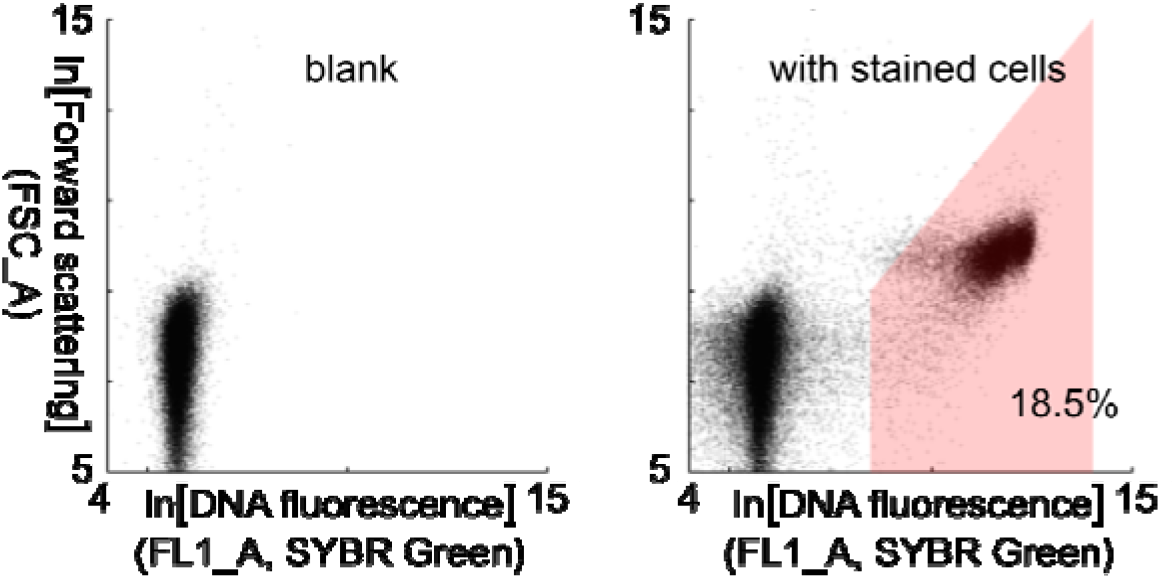
Gating for flow cytometry experiments. For all flow cytometry experiments, cells were stained with SYBR Green I dye and incubated for at least 10 minutes before measurement (see Methods). 10 μL of events were measured (scattering and fluorescence) at the slow rate (14 μL/min) and then analyzed with MATLAB R2015B. A scatter plot of the measured events are shown for a blank (left) and cell sample (right). We only focused on the forward scattering and green fluorescence dimensions. The gating used for all samples is shown by the red region. In the sample shown, the gate captured 18.5% of the events, which were taken to be the bacterial cells.

**Supplementary Figure 3:**
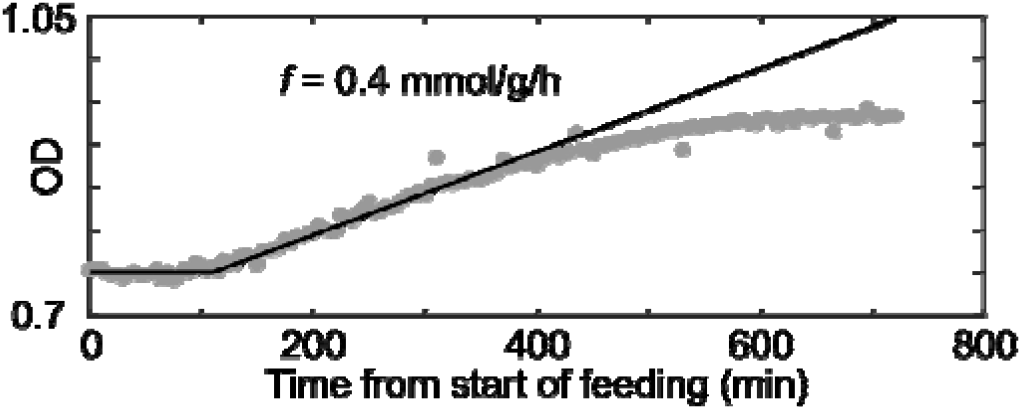
Optical density flattens beyond 6 h. For the given glucose pulsing experiment (*f* = 0.4 mmol/g/h), optical density (OD) was measured beyond the normal 6 h. Measurements are indicated by grey dots. Beyond the default experiment time of 6 h, the OD begins to flatten and ceases linear increasing. The black line indicates the predicted OD when no flattening is assumed. The prediction was based off the empirical model of Fig 2a (as described in Methods) and the calculated division yield from Fig 3.

**Supplementary Figure 4:**
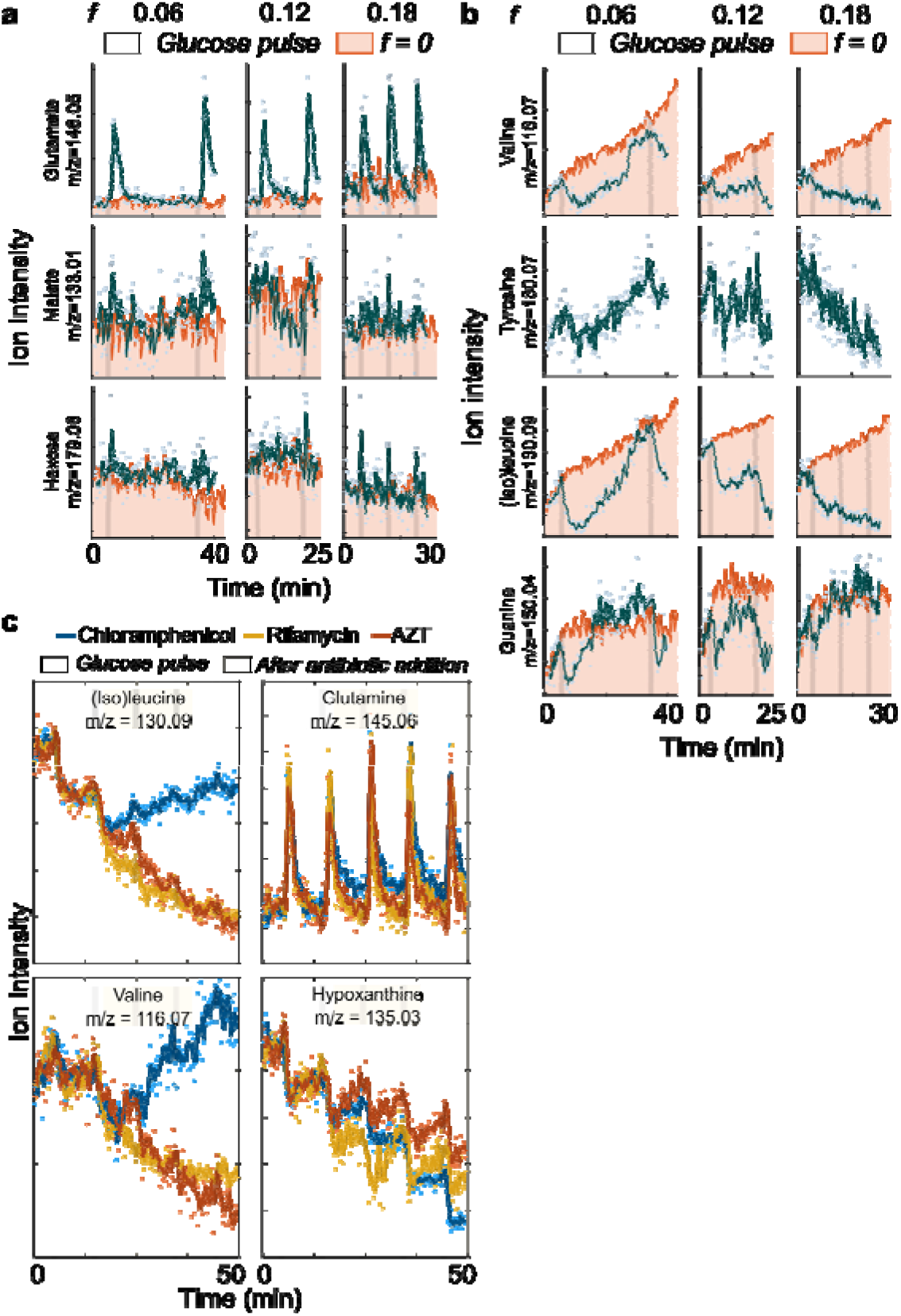
Additional real-time metabolomics data. **a**, Other central metabolites exhibited concentration spikes with glucose pulses at non-division TI feedrates (*f* = 0.06, 0.12, and 0.18 mmol/g/h). The TI feedrate is abbreviated as *f* (units: mmol glucose/g dry cell weight/hour). Glucose pulses are indicated by the grey bars, and the pink region shows a no pulse control. Dots are ion intensity measurements. Solid lines are a moving average filter of the measured ion intensity **b**, Accumulated valine, tyrosine, (iso)leucine, and guanine depleted and recovered after pulse occurrence suggesting protein and nucleic synthesis. **c**, Other amino acids and hypoxanthine were affected by corresponding antibiotics (*f* = 0.18 mmol/g/h). Antibiotics were added one minute after the second pulse (yellow region). Chloramphenicol (blue) inhibits protein biosynthesis, rifamycin (orange) inhibits RNA polymerase, and azidothymidine (AZT; red) inhibits DNA synthesis. The ion for tyrosine could not be annotated for the *f* = 0 mmol/g/h measurement.

**Supplementary Figure 5:**
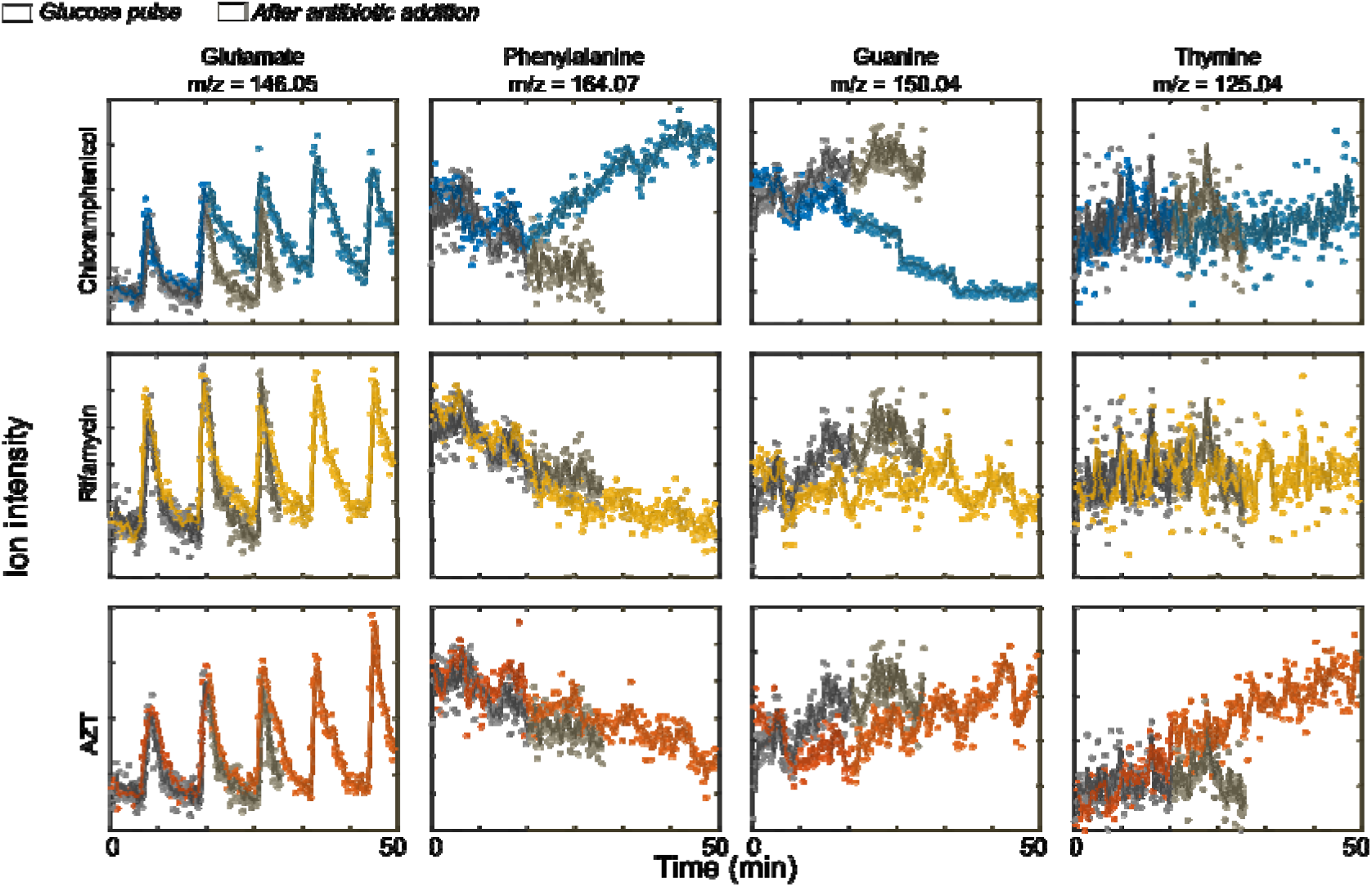
Antibiotic metabolomics data with no antibiotic control. Data from Fig 4c is plotted against the no antibiotic control condition (*f* = 0.18 mmol/g/h from Fig 4a). Black dots indicate the ion intensity for the no antibiotic condition and the solid lines indicate the moving average filter of the ion intensity.

**Supplementary Figure 6:**
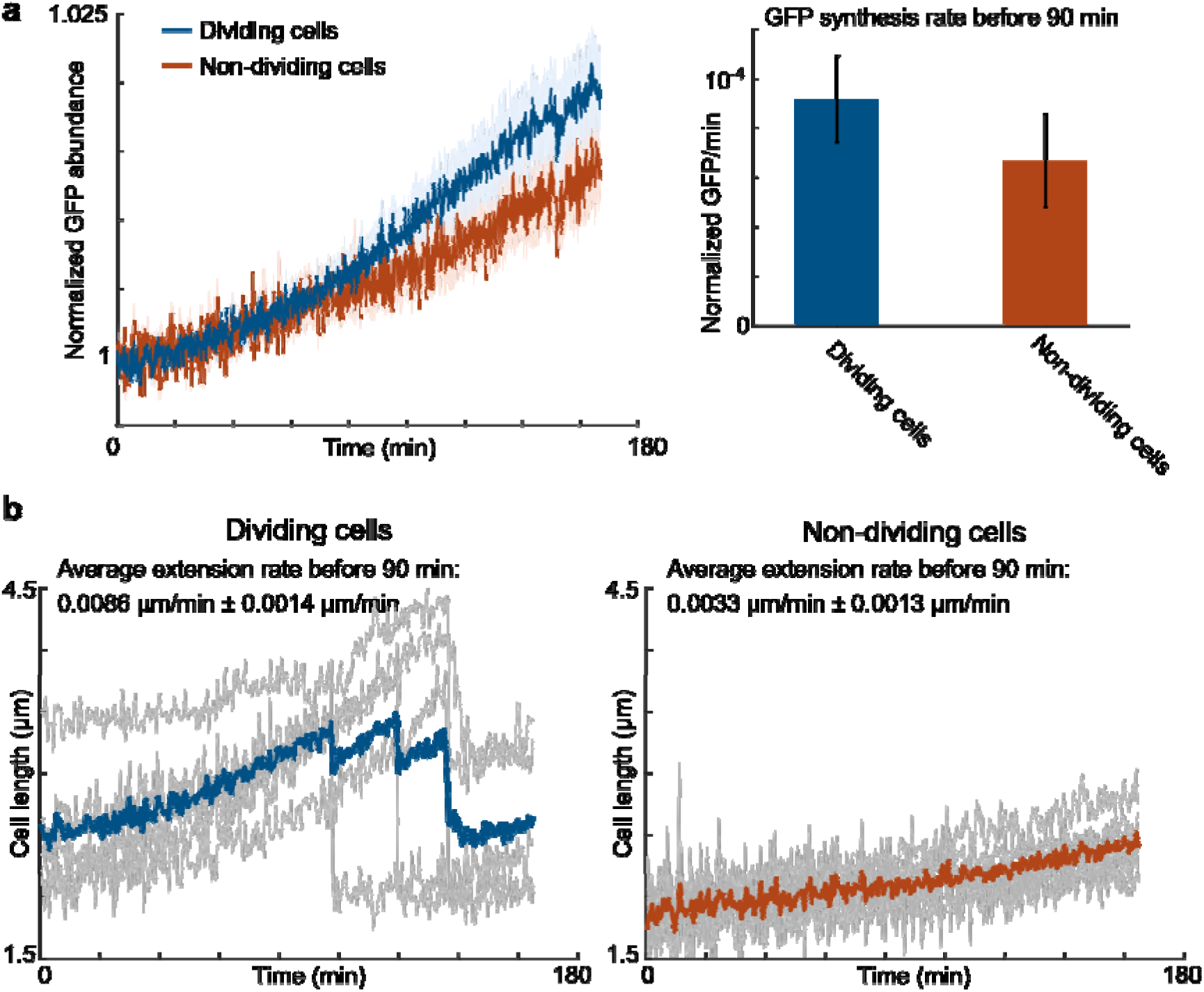
All cells make protein and increase in size with glucose feed. Cell length and GFP expression was measured using the microfluidic-based microscopy platform for glucose pulsing periods of 4 minutes. Two groups were determined: a dividing group (blue), where cells divided within the given experiment time (5 h), and a non-dividing group (red), where the cells did not divide within the 5 h. **a**, Both dividing and non-dividing cells made protein as indicated by signal from constitutive GFP expression. Solid lines indicate the moving average for each given group. Lightly shaded regions indicate average ± standard error of the cells (*n* = 5). GFP synthesis rate was calculated before division (~90 min) by using linear fitting. The bar graph shows the GFP synthesis rate for the both dividing and non-dividing cells. Values are mean ± standard error of independent cells (*n* = 5). **b**, Cell length increased with pulsing in both the dividing versus non-dividing subpopulation. Solid lines (blue and red) indicate the average of the individual cells. Grey lines indicate lengths of individual cells over time. Average cell extension rate before division was calculated using linear fitting for each cell. Values are mean ± standard error of independent cells (*n* = 5).

**Supplementary Figure 7:**
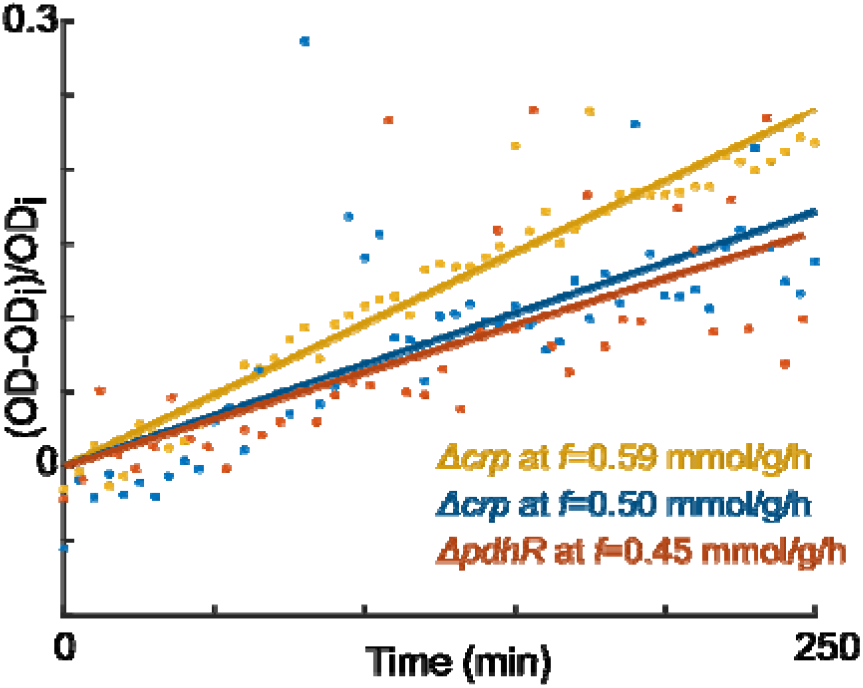
The *crp* and *pdhR* mutant strains have no lag. At normally lag-inducing TI feedrates (Fig 2a), a strain with genetic deletion of *crp* or *pdhR* showed no lag phase with glucose pulse feeding.

**Supplementary Figure 8:**
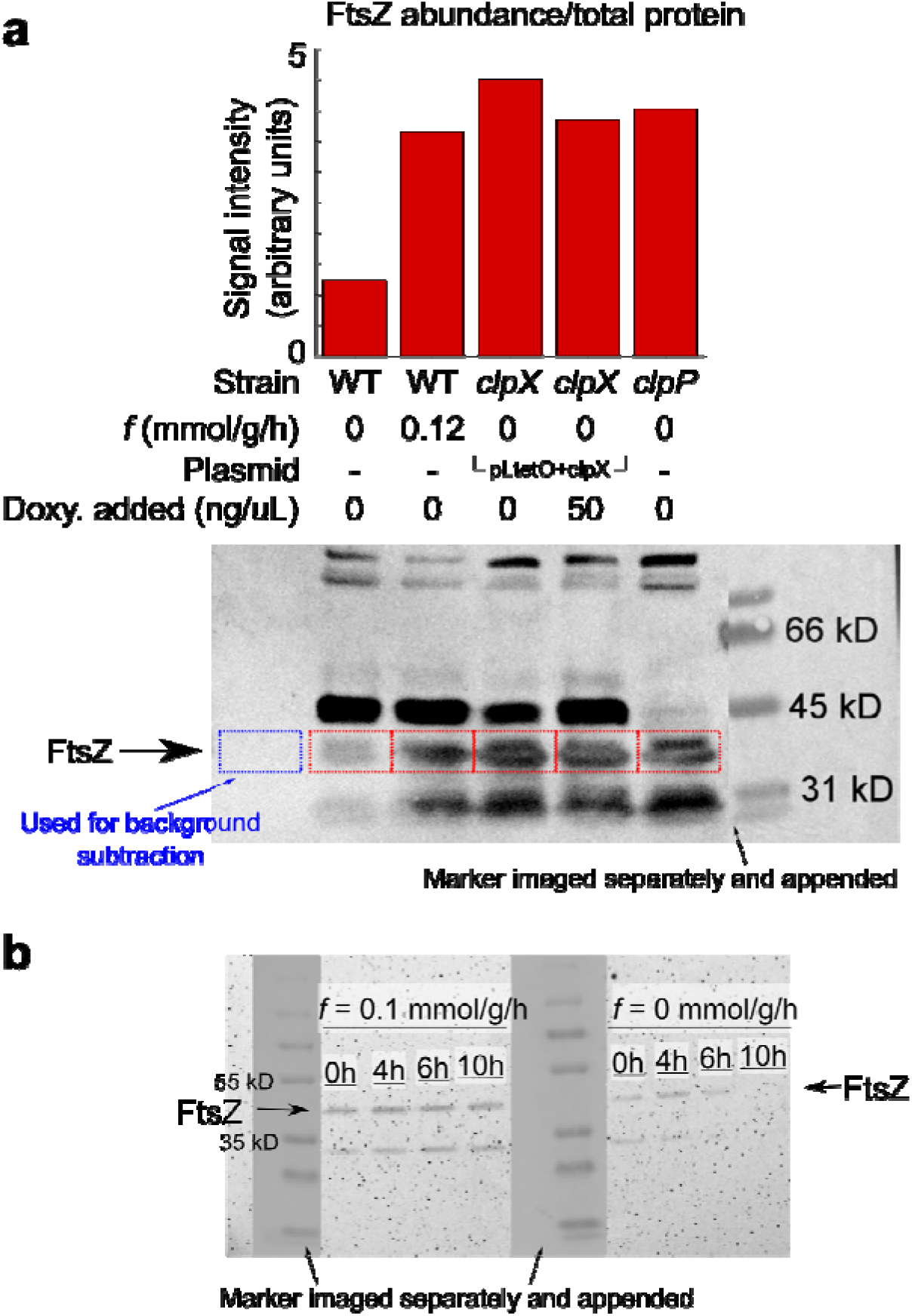
Western blots validate ClpXP-mediated degradation of FtsZ *in vivo* during starvation and synthesis of FtsZ with glucose pulsing. 1.5 ng total protein was loaded into each lane. Protein marker was imaged separately with bright-field and appended to blot with exact positioning. (**a**) After 16 h, relative FtsZ abundance (from blot directly below) is much lower in wild-type cells without any glucose pulsing (*f* = 0 mmol/g/h) compared to conditions with glucose pulsing (*f* = 0.12 mmol/g/h) or in strains absent of ClpXP machinery (*clpX* and *clpP*). Supplemental synthesis of ClpX within a *clpX* strain via expression off the pLtetO + clpX plasmid shows less FtsZ when ClpX synthesis is on (50 ng/uL doxycycline added) versus off (0 ng/uL doxycycline). Bordered areas were quantified with MATLAB 2015b. The subtracted background is indicated by the blue border. (**b**) A time course immunoblot shows depletion of FtsZ in the no pulse condition (*f* = 0 mmol/g/h) versus the pulsing condition (*f* = 0.1 mmol/g/h) across 10 h. Time indicates sampling points from the beginning of pulsing (2 hours into glucose starvation) for both experiments.

**Supplementary Figure 9:**
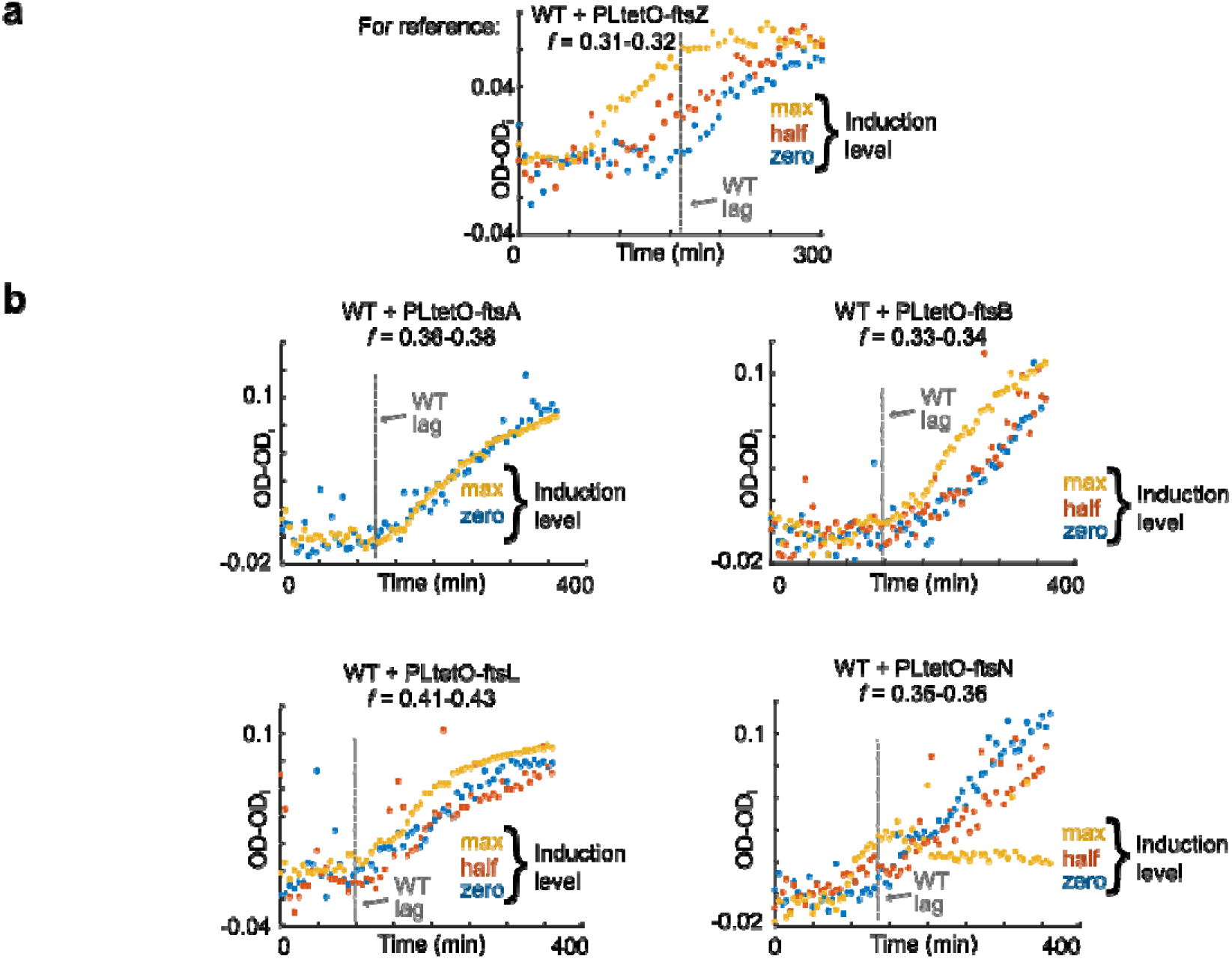
Titrations of other division proteins support FtsZ as division determinant. **a**, Fig 6f is reproduced here as reference and shows the decrease of lag time monotonic to the FtsZ induction level. Additional protein was titrated via plasmid-based, inducible expression. For induction, max, half, and zero correspond to addition of 50, 10, and 0 ng/uL of doxycycline respectively. Units of TI feedrate *f* are mmol/g/h. **b**, Lag times do not decrease with induction level of other division proteins (FtsL, FtsB, and FtsA). FtsB and FtsL induction minimally increased more division after lag end. Lag time decreases with FtsN induction, and total division is decreased as shown by the lower final OD.

**Supplementary Figure 10:**
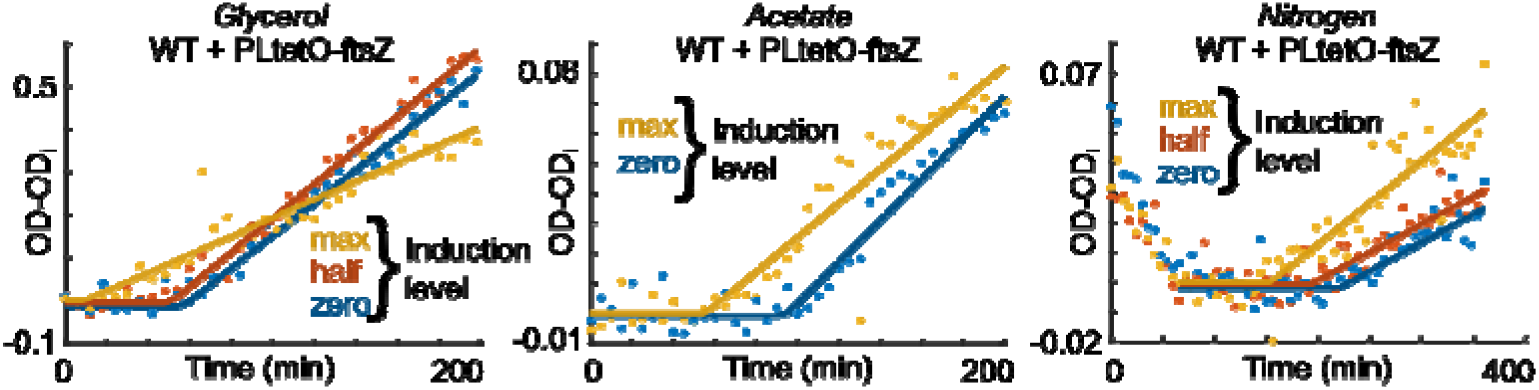
FtsZ-limited division applies for various nutrient limitations. Supplemental FtsZ titration reduced the lag time in starved cells that were pulse-fed with the limiting nutrients glycerol, acetate, or ammonium. For induction, max, half, and zero correspond to addition of 50, 10, and 0 ng/uL of doxycycline respectively. For the glycerol experiment, cells were grown in glycerol to exponential phase prior to starvation. Pulse concentration was 38 μM glycerol, and the TI feedrate was 0.6 mmol glycerol/g DCW/hour where OD 1 corresponds to 0.54 g DCW/L^52^. For the acetate experiment, cells were grown on acetate as the sole carbon source prior to starvation and pulse feeding. Pulse concentration was 88 μM sodium acetate at a TI feedrate of 3.0 mmol acetate/g DCW/hour where OD 1 corresponds to 0.44 g DCW/L^52^. For the ammonium experiment, cells were grown in M9 glucose media, then starved in media without ammonium, and consequently pulse fed. Pulse concentrations were 1.5 μM ammonium sulfate, and the TI feedrate was 0.045 mmol ammonium sulfate/g DCW/h.

**Supplementary Figure 11:**
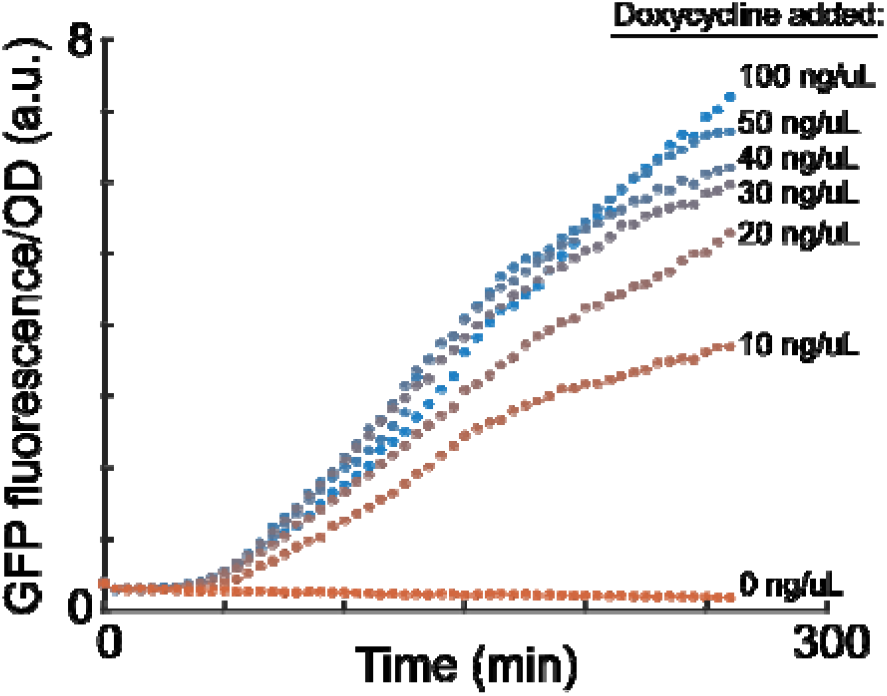
Titration curve of inducible plasmid with GFP. The parent plasmid pJKR-L-tetR^45^ was used to determine the appropriate induction level for all titration experiments. GFP expression is driven by the titratable pLtetO promoter. Normalized GFP levels are shown for different levels of inducer doxycycline over time for microplate cultivations. Time at 0 indicates the addition of doxycycline and start of plate reader measurement. For titration experiments, 50 ng/uL was selected as high, 10 ng/uL as half, and 0 ng/uL as zero expression.

**Supplementary Table 2.**
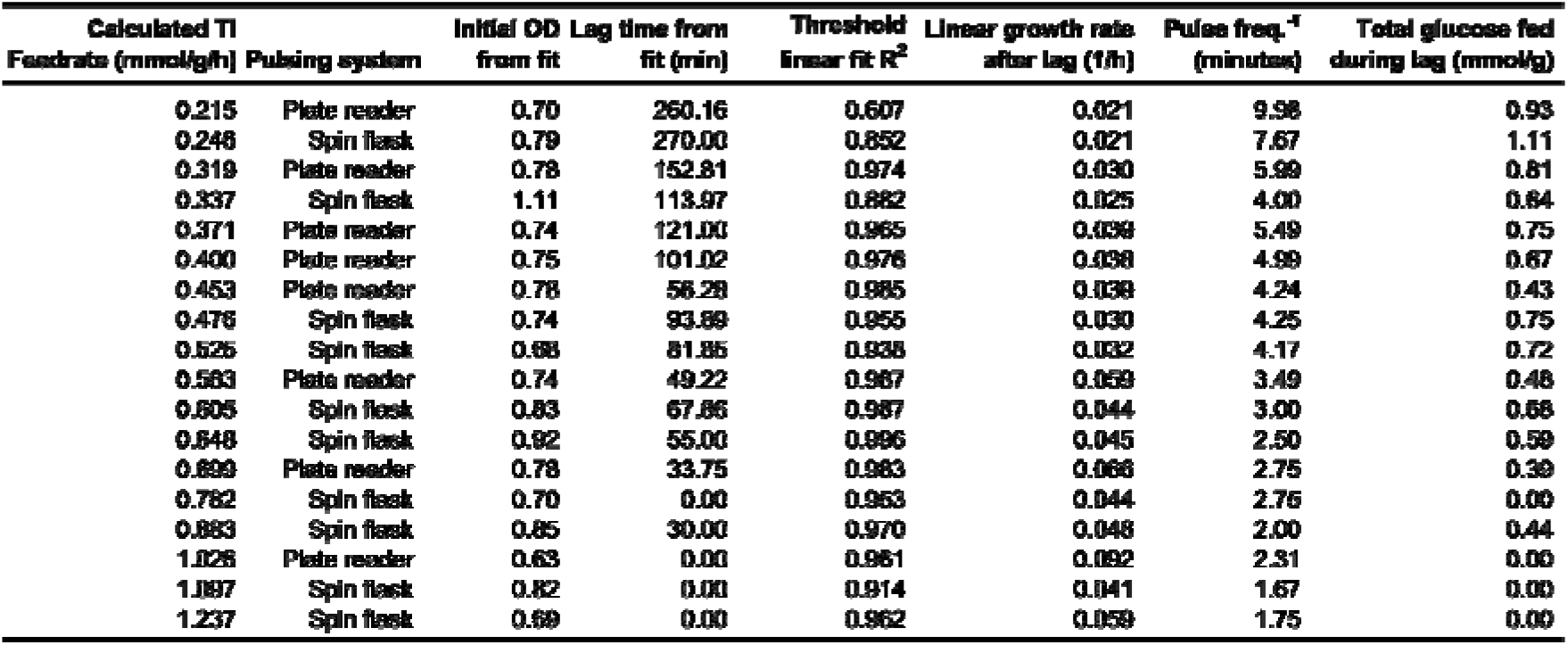
Summary information for wild-type pulse feed experiments. For the units, mmol is mmol of glucose, and g is grams dry cell weight of *E. coli*. Source data for this table is available in Supplementary Data 2.

| Calculated TI |  | Initial OD | Lag time from | Threshold | Linear growth rate | Pulse freq.<sup>-1</sup> | Total glucose fed |
| --- | --- | --- | --- | --- | --- | --- | --- |
| Feedrate (mmol/g/h) | Pulsing system | from fit | fit (min) | linear fit R<sup>2</sup> | after lag (1/h) | (minutes) | during lag (mmol/g) |
| 0.215 | Plate reader | 0.70 | 260.16 | 0.607 | 0.021 | 9.98 | 0.93 |
| 0.246 | Spin flask | 0.79 | 270.00 | 0.852 | 0.021 | 7.67 | 1.11 |
| 0.319 | Plate reader | 0.78 | 152.81 | 0.974 | 0.030 | 5.99 | 0.81 |
| 0.337 | Spin flask | 1.11 | 113.97 | 0.882 | 0.025 | 4.00 | 0.64 |
| 0.371 | Plate reader | 0.74 | 121.00 | 0.965 | 0.039 | 5.49 | 0.75 |
| 0.400 | Plate reader | 0.75 | 101.02 | 0.976 | 0.038 | 4.99 | 0.67 |
| 0.453 | Plate reader | 0.78 | 56.28 | 0.985 | 0.038 | 4.24 | 0.43 |
| 0.476 | Spin flask | 0.74 | 93.89 | 0.955 | 0.030 | 4.25 | 0.75 |
| 0.526 | Spin flask | 0.68 | 81.85 | 0.938 | 0.032 | 4.17 | 0.72 |
| 0.563 | Plate reader | 0.74 | 49.22 | 0.967 | 0.039 | 3.49 | 0.48 |
| 0.605 | Spin flask | 0.83 | 67.86 | 0.987 | 0.044 | 3.00 | 0.68 |
| 0.648 | Spin flask | 0.92 | 55.00 | 0.996 | 0.045 | 2.50 | 0.59 |
| 0.699 | Plate reader | 0.78 | 33.75 | 0.983 | 0.066 | 2.75 | 0.39 |
| 0.782 | Spin flask | 0.70 | 0.00 | 0.963 | 0.044 | 2.76 | 0.00 |
| 0.883 | Spin flask | 0.85 | 30.00 | 0.970 | 0.048 | 2.00 | 0.44 |
| 1.026 | Plate reader | 0.63 | 0.00 | 0.961 | 0.092 | 2.31 | 0.00 |
| 1.187 | Spin flask | 0.82 | 0.00 | 0.914 | 0.041 | 1.67 | 0.00 |
| 1.237 | Spin flask | 0.69 | 0.00 | 0.962 | 0.059 | 1.75 | 0.00 |

**Supplementary Table 3.** C^13^ Labeled fraction of measured amino acids and glycogen in washed, hydrolyzed extract. All measurements are after 6 hours of pulsing. Values are the mean ± standard error of independent biological replicates (*n* = 3 for amino acid samples, *n* = 2 for glycogen). ^+^Class designation is from ^17^

| Compound [% of native carbons in fragment] | Fraction of [+X] fragment (n = 3, biological) at given TI feedrate |  |  |  |
| --- | --- | --- | --- | --- |
|  | 0 mmol/g/h | 0.06 mmol/g/h | 0.12 mmol/g/h | 0.18 mmol/g/h |
| **Class 1<sup>+</sup> - Non-accumulating amino acids** |  |  |  |  |
| Alanine [3] | 0.002 ± 0.002 | 0.025 ± 0.003 | 0.058 ± 0.007 | 0.060 ± 0.003 |
| Aspartate [4] | 0 ± 0 | 0.014 ± 0.001 | 0.038 ± 0.001 | 0.060 ± 0.003 |
| Glutamate [5] | 0.0006 ± 0.0001 | 0.017 ± 0.001 | 0.039 ± 0.001 | 0.065 ± 0.002 |
| Glycine [2] | 0.0013 ± 0.0007 | 0.022 ± 0.001 | 0.04 ± 0.01 | 0.070 ± 0.003 |
| Proline [4] | 0.0005 ± 0.0003 | 0.0061 ± 0.0002 | 0.013 ± 0.001 | 0.018 ± 0.001 |
| Serine [2] | 0.001 ± 0.001 | 0.026 ± 0.002 | 0.039 ± 0.006 | 0.057 ± 0.002 |
| Threonine [4] | 0 ± 0 | 0.008 ± 0.001 | 0.022 ± 0.002 | 0.039 ± 0.004 |
| **Class 2<sup>+</sup> - Accumulating amino acids** |  |  |  |  |
| Isoleucine [5] | 0.003 ± 0.001 | 0.003 ± 0.001 | 0.006 ± 0.003 | 0.00013 ± 0.00001 |
| Leucine [5] | [+6] not detected |  |  |  |
| Lysine [5] | [+5] not detected |  |  |  |
| Methionine [4] | [+4] not detected |  |  |  |
| Phenylalanine [8] | [+8] not detected |  |  |  |
| Tyrosine [9] | [+9] not detected |  |  |  |
| Valine [4] | 0.002 ± 0.001 | 0.0010 ± 0.0002 | 0.002 ± 0.001 | 0.008 ± 0.002 |
| Glycogen [5] | 0.0004 ± 0.0001 | 0.00028 ± 0.00002 | 0.0007 ± 0.0003 |  |

## Supplementary Information

### • PDF files

#### ∘ Supplementary Information

FtsZ model development, parametrization, and analytical solution. Supplementary Figures 12-14. Supplementary Tables 4-6. Supplementary references.

### • Video files

#### ∘ Video 1

6 cells recorded during microfluidic single-cell microscopy (pulse every 4 minutes).

### • ZIP files

#### ∘ Supplementary Data 1

GenBank files for all plasmids used in this study.

#### ∘ Supplementary Data 2

All code and data used for analysis and figures.

### • Excel files

#### ∘ Supplementary Table 1

Annotation and ion counts for all metabolomics data.

